# An epigenetically distinct HSC subset supports thymic reconstitution

**DOI:** 10.1101/2024.06.06.597775

**Authors:** Harold K. Elias, Sneha Mitra, Marina B. da Silva, Adhithi Rajagopalan, Brianna Gipson, Nicole Lee, Anastasia I. Kousa, Mohamed A.E. Ali, Simon Grassman, Xiaoqun Zhang, Susan DeWolf, Melody Smith, Hana Andrlova, Kimon V. Argyropoulos, Roshan Sharma, Teng Fei, Joseph C Sun, Cynthia E. Dunbar, Christopher Y Park, Christina S. Leslie, Avinash Bhandoola, Marcel R.M. van den Brink

**Affiliations:** Department of Immunology, Sloan Kettering Institute, Memorial Sloan Kettering Cancer Center, New York, NY, 10065, USA; National Institutes of Health (NIH), Bethesda, MD, 20892, USA, USA; Computational and Systems Biology, Sloan Kettering Institute, Memorial Sloan Kettering Cancer Center, New York, NY, 10065, USA; Department of Pathology, New York University Grossman School of Medicine, New York, NY, 10016, USA; Division of Blood and Marrow Transplantation and Cellular Therapy, Department of Medicine, Stanford University School of Medicine, Stanford, CA, 94305; Leukemia Service, Department of Medicine, Memorial Sloan Kettering Cancer Center, New York, NY, 10065, USA; Single-cell Analytics and Innovation Lab, Sloan Kettering Institute, Memorial Sloan Kettering Cancer Center, New York, NY, 10065, USA; Department of Epidemiology and Biostatistics, Memorial Sloan Kettering Cancer Center, New York, NY, 10065, USA; Translational Stem Cell Biology Branch, National Heart, Lung, and Blood Institute, NIH, Bethesda, MD, 20892, USA; Laboratory of Genome Integrity, Center for Cancer Research, National Cancer Institute, NIH, Bethesda, MD, 20892, USA; Department of Immunology and Microbial Pathogenesis, Weill Cornell Medical College; New York, NY; Department of Medicine, Memorial Sloan Kettering Cancer Center, New York, NY, 10065, USA

**Author notes:** Correspondence (HKE); (MRMvdB). These authors contributed equally (Marina B. da Silva, Adhithi Rajagopalan).

**Keywords:** Hematopoietic stem cells, lymphoid differentiation, epigenetic priming, thymus, immune reconstitution, immunosenescence

## Abstract

Hematopoietic stem cells (HSCs) with multilineage potential are critical for effective T cell reconstitution and restoration of the adaptive immune system after allogeneic Hematopoietic Cell Transplantation (allo-HCT). The Kit^lo^ subset of HSCs is enriched for multipotential precursors,^1, 2^ but their T-cell lineage potential has not been well-characterized. We therefore studied the thymic reconstituting and T-cell potential of Kit^lo^ HSCs. Using a preclinical allo-HCT model, we demonstrate that Kit^lo^ HSCs support better thymic recovery, and T-cell reconstitution resulting in improved T cell responses to infection post-HCT. Furthermore, Kit^lo^ HSCs with augmented BM lymphopoiesis mitigate age-associated thymic alterations, thus enhancing T-cell recovery in middle-aged hosts. We find the frequency of the Kit^lo^ subset declines with age, providing one explanation for the reduced frequency of T-competent HSCs and reduced T-lymphopoietic potential in BM precursors of aged mice.^3, 4, 5^ Chromatin profiling revealed that Kit^lo^ HSCs exhibit higher activity of lymphoid-specifying transcription factors (TFs), including *Zbtb1*. Deletion of *Zbtb1* in Kit^lo^ HSCs diminished their T-cell potential, while reinstating *Zbtb1* in megakaryocytic-biased Kit^hi^ HSCs rescued T-cell potential, *in vitro* and *in vivo*. Finally, we discover an analogous Kit^lo^ HSC subset with enhanced lymphoid potential in human bone marrow. Our results demonstrate that Kit^lo^ HSCs with enhanced lymphoid potential have a distinct underlying epigenetic program.

## INTRODUCTION

T lymphocytes play a crucial role in the adaptive immune response. A highly diverse T cell pool is required to recognize and eliminate foreign pathogens while also maintaining self-tolerance. The thymus continuously produces T cells, but with advancing age, its regenerative capacity declines,^6^ leading to decreased thymic output and compromised T cell diversity. Furthermore, the thymus is also vulnerable to acute damage from infections, cancer therapies, and conditioning regimens for allogeneic hematopoietic cell transplantation (allo-HCT), which results in prolonged T cell lymphopenia, thereby increasing the risk of infections and cancer relapse, contributing to transplant-related complications and mortality.^7^ Likely for these reasons, early T-cell reconstitution is a positive prognostic indicator of allo-HCT outcomes.^8, 9^

Post-transplant T cell reconstitution requires a steady supply of hematopoietic stem cells (HSCs)-derived lymphoid progenitors^10^ as well as thymic recovery. Prior research has demonstrated that thymic integrity is dependent on thymocyte-stromal crosstalk, particularly in the thymic epithelial cell (TEC) compartment.^11, 12, 13^ Age-associated reduced lymphoid potential originating at the level of HSCs,^14^ coupled with thymic decline^6^ contribute to delayed immune recovery. Therefore, identifying strategies to augment T lymphopoiesis and promote thymic regeneration is an unmet clinical need.

Single-cell transplantation with tracking and sequencing of progeny^15, 16, 17, 18, 19, 20^ has uncovered heterogeneity in self renewal capacity as well as lineage output among reconstituting HSCs-highlighting HSCs with multilineage versus lineage-biased potential. These studies showed that, while only a fraction of HSCs generate “balanced” multilineage output, the majority exhibit a diverse range of lineage potential following transplantation-varying in their contributions and reconstitution kinetics for each lineage, thereby showcasing their inherent biases. These lineage biases are intrinsically stable and maintained following transplantation.^21, 22, 23^ At the molecular level, these cell autonomous features are mediated by epigenetic configuration and are divergent from their transcriptional state.^18, 24^ Although this suggests the existence of a highly organized and predictable framework for lineage-restricted fates of long-term self-renewing HSCs, the molecular identities, gene regulatory networks and functional implications of these differences governing lymphoid fate decisions remain largely unexplored.

Kit^lo^ HSCs have previously been described to exhibit better self-renewal and multipotency, including improved T-cell reconstitution in syngeneic transplant models. ^1, 2^However, their thymic reconstituting ability, age-related changes, as well as the molecular basis for their enhanced T-cell lymphoid potential are unknown. Here, we delineate the pivotal role of Kit^lo^ HSCs in orchestrating thymic regeneration and immune reconstitution in both young and aged hosts. We find the frequency of the Kit^lo^ subset declines with age. Leveraging integrated single cell transcriptional and epigenetic profiling, we uncover the distinct epigenetic signature of Kit^lo^ HSCs, highlighting the presence of lymphoid-specifying transcription factor *Zbtb1* and its role in enhancing T-cell lymphoid potential. Additionally, we identify an analogous HSC subset in human bone marrow and demonstrate their enhanced lymphoid potential.

## RESULTS

### Reduced lymphoid output in aged HSCs impairs thymic recovery

Previous studies have shown impaired lymphoid reconstitution with aged HSCs in a syngeneic transplant model, yet this has not been explored using an allogeneic setting. We used a pre-clinical allo-HCT model to analyze immune reconstitution of young and old donors. To evaluate if HSCs with differences in lymphoid progenitor production could impact thymic recovery, we transplanted Lin−Sca-1+Kit+ (LSK) CD34−CD150+CD48− cells LT-HSCs from young adult (2-mo) or old (24-mo) C57BL/6 mice with rescue BM cells (CD45.1) into lethally irradiated young BALB/cJ recipients (**Figure S1A**). Because HSC-derived thymic progeny emerge within 5-6 weeks,^25^ we established an 8-week harvest timepoint. This interval allowed sufficient time for HSC-derived progenitors to repopulate the thymus, thereby providing a better understanding of thymic regeneration dynamics and the efficacy of HSC transplantation in promoting thymic reconstitution.

Consistent with prior reports, eight weeks post-HCT, we found that young HSCs supported balanced lineage reconstitution, while old HSCs exhibited significantly impaired T-cell reconstitution in the peripheral blood (PB), despite comparable reconstitution patterns at four weeks (**Figures S1B-S1E**). Furthermore, we observed greater multilineage potential of young HSCs, including lymphoid and myeloid reconstitution (LMPP/MPP4 and CLP, **Figures S1F and S1G**), when compared to old HSCs. Furthermore, young HSC recipients had significantly greater thymic cellularity and reconstitution of precursor and mature thymocytes than recipients of old HSCs (**Figures S1H-S1J**). In conclusion, old HSCs have significantly less multi-lineage potential, but especially T lymphoid potential than young HSCs in young allo-HCT recipients.

### Age-related decline in multilineage HSCs marked by low Kit expression

Recent reports have shown remarkable functional heterogeneity of HSC^26, 27, 28, 29, 30, 31^ subsets, highlighting discordance between current HSC phenotypic definitions to their molecular identities.^16, 17, 18^ To identify HSCs with an enriched multilineage program and gain insights into their transcriptional and epigenetic framework, we performed multiome single-cell RNA and ATAC sequencing on hematopoietic stem cells (LT-HSCs: Lineage^-^CD34^-^CD48^-^CD150^+^Sca-1^+^c-Kit^+^) from 2-mo (young) or 24-mo (old) female mice.

To define HSC subsets, we performed unsupervised Leiden cluster analysis on the multiome scRNA-seq data from our young cohort (**Figure S2A-S2B**). Using nomenclature and signatures derived from previous studies, ^15, 16, 18, 32, 33, 34, 35^ we mapped all unsupervised clusters into five main subsets (**Figure S2C**): Quiescent HSC (q-HSC), Platelet-biased HSC (Mgk-HSC), Multilineage HSC (MLin-HSC), Proliferative HSC (p-HSC), and a novel Intermediate HSC (Int-HSC) that showed overlapping features with Mgk-HSC, MLin-HSC and p-HSC subsets (**Data Table 1, Figure S2D**). To identify HSCs with multilineage potential, we compared multilineage enriched MLin-HSC versus low output q-HSC subsets in young HSCs and queried for previously described bonafide HSC markers (*Hlf, Kit, Neo1, Procr, CD244a, Pdzk1ip1, Ly9, Hoxb5*). *Hlf* and *Kit* had significantly higher expression in q-HSCs compared to MLin-HSCs (**Figure 1A**). *Hlf* has been previously implicated in HSC quiescence,^36, 37, 38^ whereas *Kit* enriches for platelet-biased HSCs.^2^ To further analyze the composition of HSCs with varying Kit expression, we defined Kit high and low HSC populations (Kit^hi^ and Kit^lo^ HSCs) based on the top and bottom 20% of *Kit* gene expression levels. We observed lower Kit expression in MLin-HSCs (**Figure 1B**) and increased representation of q-HSC and MLin-HSCs within the Kit^lo^ HSCs (**Figures 1C and S2E**), consistent with prior reports of increased self-renewal capacity and multipotency in Kit^lo^ HSCs.^1, 2^

**Figure 1:**
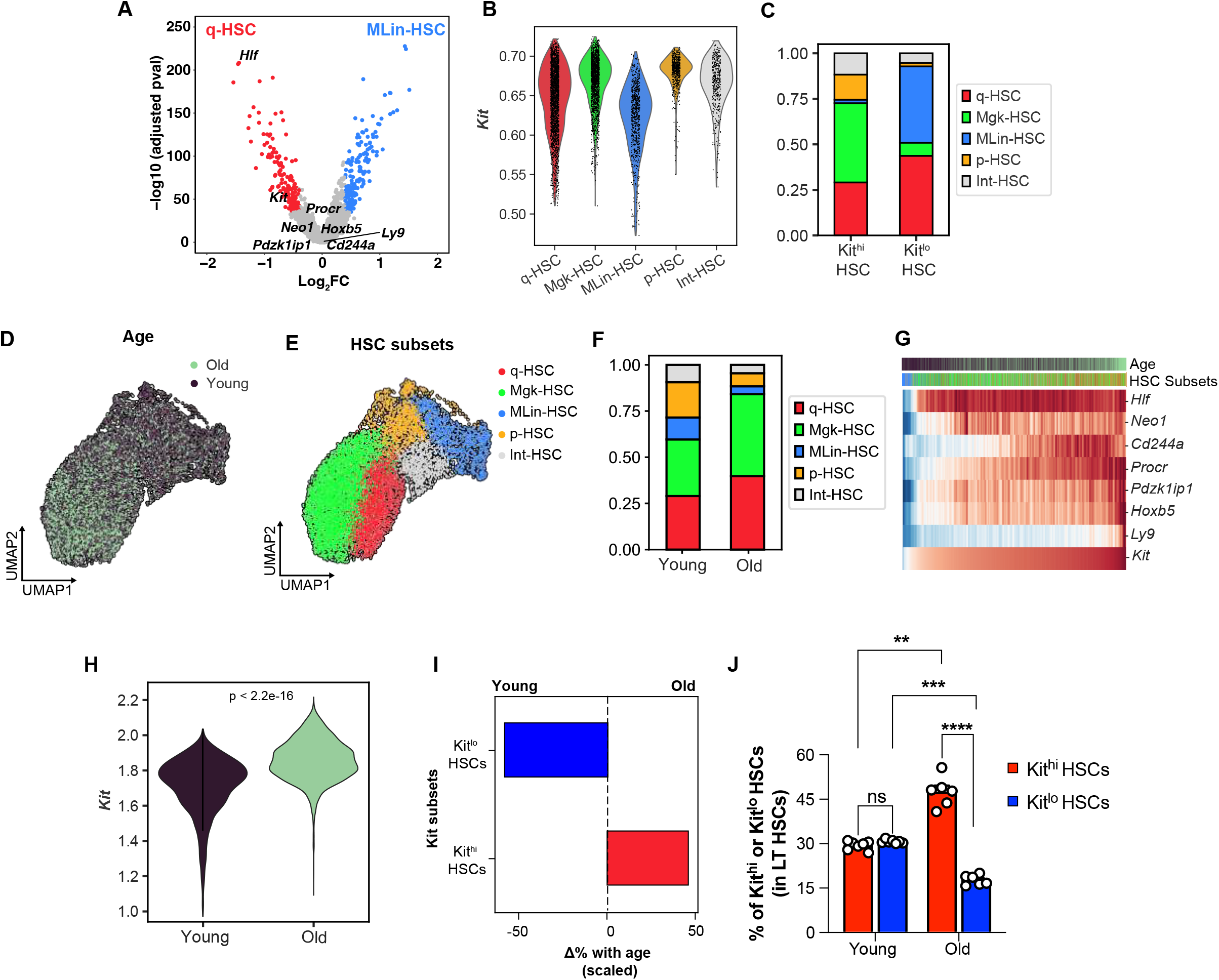
Age-related decline in multilineage HSCs marked by low Kit expression. (**A-H**) Multiome single-cell RNA and ATAC sequencing was performed on HSC (Lineage-Sca-1+cKit+ CD34-CD48-Flt3-CD150+) cells isolated from 2-mo (young) or 24-mo (old) female C57BL/6 mice. nTOTAL =12,350 cells. (**A**) Volcano plot of DGE analysis between MLin-HSC vs. q-HSC subsets in young HSCs, highlighting bonafide HSC markers. (**B**) Violin plot for imputed *Kit* gene expression by annotated HSC subsets (as described in **Fig. S2C**. (**C**) Frequency of computationally defined HSC subtypes in Kit^hi^ and Kit^lo^ subsets in young HSCs. (**D-E**) Combined Uniform manifold approximation and projection (UMAP) of young and old HSCs after Harmony batch correction annotated by age **(D)** and annotated HSC subsets (q-HSC: Quiescent HSCs; Mgk-HSCs: Platelet-biased HSCs; Mlin-HSC: Multilineage HSCs; p-HSC: Proliferative HSCs; Int-HSC: Intermediate HSCs) **(E)**. **(F)** Frequency of HSC subtypes in young and old HSCs. **(G)** Heatmap showing gene expression of bonafide HSC markers ordered by increasing Kit expression. **(H)** Violin plot for imputed Kit gene expression by age. Statistical analysis was performed using the Wilcoxon test. **(I)** Scaled change in frequency for each Kit HSC subset with age. **(J)** FACS analysis showing frequency of Kit^hi^ and Kit^lo^ HSCs by age. All data are from n=6 mice/group. Error bars represent mean ± SEM. **P<0.01, ***P<0.001. P values calculated by two-way ANOVA.

To assess for age-associated changes, we first compared the transcriptional profile of young and old HSCs. In our differential gene analysis, we found that HSCs from young mice had higher expression of genes related to cell cycle (*Ccnd3*, *Cdc25a*, *Cdk1)*, and cell polarization (*Arhgap15*); conversely, old HSCs were enriched for expression of genes related to quiescence (*Egr1*, *Junb, Jun*, *Jund*, *Fos*), platelet differentiation (*vWf*, *Clu*, *Itgb3, Fhl1 and Tgm2*), and myeloid differentiation (*Selp, Neo1*) (**Data Table 2**). Next to investigate age-related shifts in HSC cluster composition, we employed *ingest()* from scanpy to query the old HSC transcriptomic data using the “trained” young reference data for HSC subsets (**Figure S2F**). Subsequently, we generated a UMAP of our young and old datasets (**Figures S2G and 1D**) to visualize the post-ingestion results (**Figures 1E**). While all HSC subsets were present in young and old cohorts (**Figure 1F**), we notably observed an increased representation of MLin-HSCs, p-HSCs, and Int-HSCs in young HSCs. Conversely q-HSCs and Mgk-HSCs were predominantly in old HSCs (**Figures S2H-S2J**). Consistent with prior work,^18, 27, 39^ these findings suggest shifts in the composition of HSC subsets with aging, indicating a reduction in multilineage potential accompanied by an increase in platelet bias.

Given the increased frequency of Mgk-HSCs in old HSCs and prior work^2^ demonstrating that Kit^hi^ HSCs are platelet-biased HSCs (**Figures S2F-S2I**), we hypothesized this could be attributed to an expansion in Kit^hi^ HSCs. We observed significantly higher *Kit* expression (**Figures 1G-1H and S2K**) in old HSCs, with an age-dependent reduction in the proportion of Kit^lo^ HSCs accompanied by an increase in the frequency of Kit^hi^ HSCs (**Figure 1I**). Flow cytometry analysis of young and old bone marrow confirmed a significant expansion of Kit^hi^ HSCs, accompanied by a concomitant decrease in Kit^lo^ HSCs in old HSCs (**Figures S2L and 1J**), consistent with our prior report.^40^ Together, these findings indicate that the frequency of multilineage Kit^lo^ HSCs declines with age.

### Young Kit^lo^ HSCs exhibit enhanced T-cell potential

Next, we sought to determine the lymphoid potential of the HSC Kit subsets. To this end, we used the S17 lymphoid progenitor assay^41^ to assess for lymphoid progenitor output of purified Kit^hi^ and Kit^lo^ HSCs from young mice (**Figure 2A**). Prior studies have elucidated the existence of functional heterogeneity among phenotypic Common Lymphoid Progenitors (CLPs) characterized by Ly6D expression,^42^ resulting in the identification of two distinct subsets: All Lymphocyte Progenitors (ALPs), also referred to as functional CLPs, and B-cell biased Lymphocyte Progenitors (BLPs). Following 12-day co-culture, we observed that young Kit^lo^ HSCs generated more CLPs and ALPs than Kit^hi^ HSCs (**Figure 2B**). To further examine their T-cell differentiation potential, we used murine artificial thymic organoids (mATO, BM-ATO)^43, 44,45^ (**Figure 2A**). Following an 8-week culture, we noted a substantial increase in overall ATO output from young Kitl° HSCs in comparison to Kit^hi^ HSCs (**Figure 2C**).

**Figure 2:**
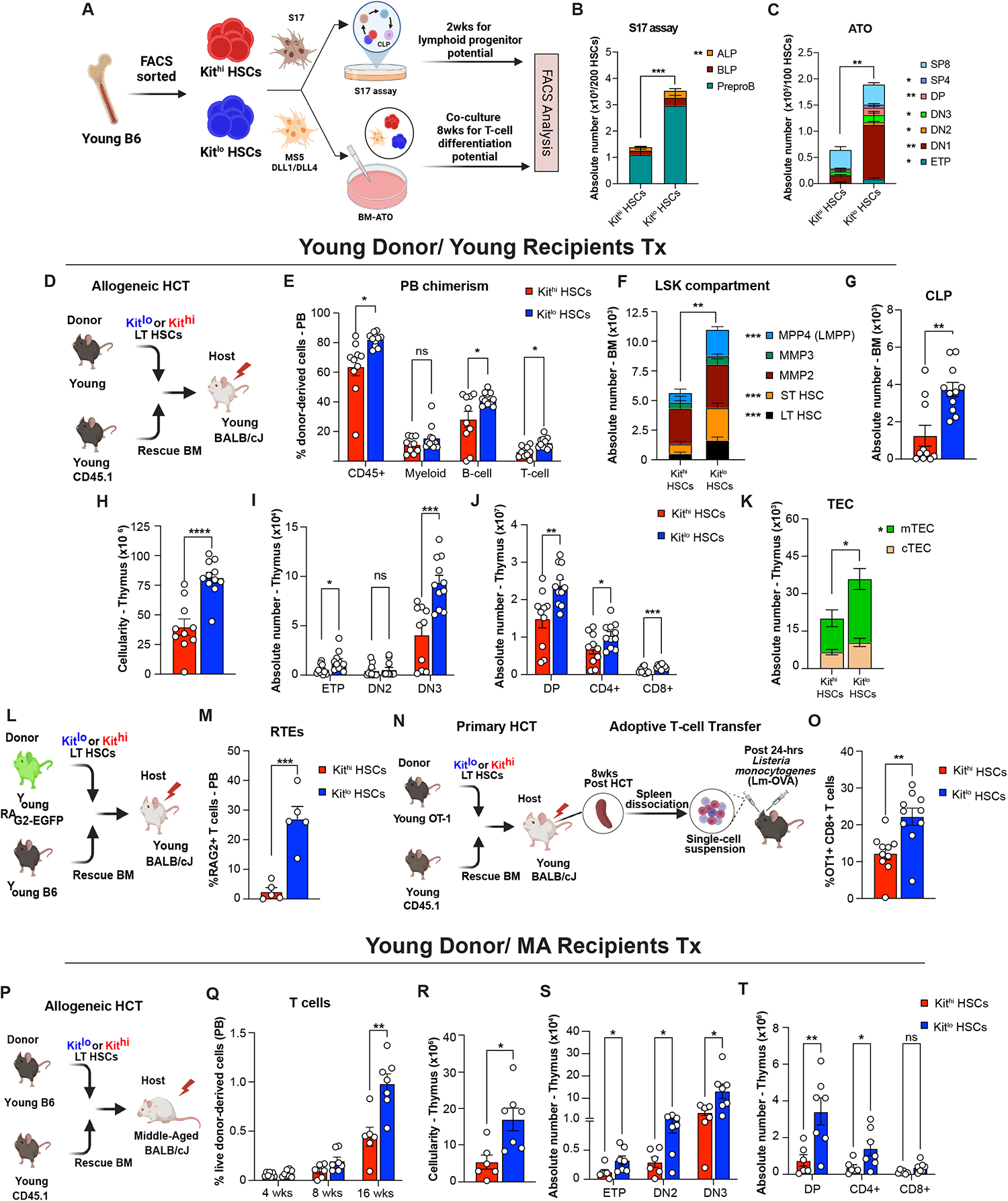
Young Kit^lo^ HSCs exhibit enhanced T-cell potential. **(A)** Experimental schema to evaluate *in vitro* lymphoid progenitor and T-cell differentiation potential of Kit^hi^ (red) and Kit^lo^ HSCs (blue) from 2-mo (young) C57BL/6 mice using the S17 lymphoid assay and murine artificial thymic organoid (mATO) respectively. Enumeration of absolute number of lymphoid progenitor cells following 14 days in S17 lymphoid assay (B) and absolute number of T-cell subsets following 8-weeks of culture in mATOs **(C).** Aggregated data across 4-5 independent experiments. **(D)** Experimental schema for competitive allogeneic HCT (allo-HCT) using Kit^hi^ (red) and Kit^lo^ HSCs (blue) from 2-mo (young) C57BL/6 mice with competitor bone marrow (BM) cells from B6.SJL-PtprcaPepcb/BoyJ mice transplanted into lethally irradiated 7-week-old (young) BALB/cJ recipients. **(E-K)** Eight weeks after competitive HCT, **(E)** frequency of donor-derived chimerism (CD45.2/ H-2Kb) of mature lineages in the peripheral blood (PB, CD45+, Myeloid cell: Gr1+CD11b+; B-cell: B220+; T-cell: CD3+), (**F-G)** Enumeration of absolute number of donor-derived cells in the BM for LSK cells (LT HSC: Lineage^-^Sca-1^+^cKit^+^ CD34^-^CD48^-^Flt3^-^CD150^+^; ST HSC: Lineage^-^Sca-1^+^cKit^+^ CD48^-^Flt3^-^ CD150^-^; MPP2: Lineage^-^Sca-1^+^cKit^+^ CD48^+^Flt3^-^CD150^-^; MPP3: Lineage^-^Sca-1^+^cKit^+^ CD48^+^Flt3^-^CD150^+^; MPP4/ LMPP: Lineage^-^Sca-1^+^cKit^+^ Flt3^+^CD150^-^) **(F),** Common Lymphoid Progenitors cells (CLP: Lineage^-^IL7Ra^+^Flt3^+^Sca^mid/lo^ Kit^lo^) **(G)**. **(H-K)** Post-HCT thymi analysis for total thymic cellularity **(H)**, **(I-J)** enumeration of absolute number of donor-derived cells for T-cell precursors (ETP: Lineage^-^ CD4^-^ CD8^-^CD44^+^ CD25^-^Kit^+^;DN2: Lineage^-^ CD4^-^ CD8^-^CD44^+^ CD25^+^; DN3: Lineage^-^ CD4^-^ CD8^-^CD44^-^ CD25^+^) **(I)**, Mature T-cells (DP: Lineage^-^ CD4^+^ CD8^+^; SP4: Lineage^-^ CD4^+^ CD8^-^; SP8: Lineage^-^ CD4^-^ CD8^+^**) (J),** and analysis of thymic CD45-compartment for Thymic Epithelial Cells (TEC: CD45^-^ EpCAM^+^), Cortical TEC (cTEC: CD45^-^ EpCAM^+^UEA-1^lo^ 6C3^hi^ MHCII^hi/lo^), and Medullary TEC (mTEC: CD45^-^ EpCAM^+^UEA-1^hi^ 6C3^lo^ MHCII^hi/lo-^) (**K**). All data are from n=10-11 mice/group. **(L)** Experimental schema to evaluate thymic function following competitive allo-HCT using young HSCs. **(M)** Frequency of donor-derived Recent Thymic Emigrants (RTE: CD3+GFP+) in the PB at 8 weeks post-HCT. All data are from n=5-6 mice/group. **(N)** Experimental schema to investigate functional response of differential Kit-expressing HSC-derived T-cells in secondary recipients to *L. monocytogenes* expressing chicken ovalbumin (LM-OVA) infection. **(O)** Spleen analysis of secondary recipients for frequency of OT1-derived chimerism of CD8+ T cells (OT1+CD8+: Va2^+^Vb5^+^H2-K^b^CD8^+^) 7 days post-infection. All data are from n=9-10 mice/group. **(P)** Experimental schema for competitive allogeneic HCT (allo-HCT) using Kit^hi^ (red) and Kit^lo^ HSCs (blue) from 2-mo (young) C57BL/6 mice with competitor bone marrow (BM) cells from B6.SJL-PtprcaPepcb/BoyJ mice transplanted into lethally irradiated 14-mo (Middle-aged) BALB/cJ recipients. **(Q)** Frequency of donor-derived T-cell chimerism at the indicated time-points. **(R-T)** Sixteen weeks after competitive HCT enumeration of total thymic cellularity **(R)**, donor-derived T-cell precursor thymocyte subsets as defined in Figure 3F **(S)**, donor-derived mature T-cell thymocyte subsets as defined in Figure 3G **(T).** All data are from n=6-7 mice/group. Error bars represent mean ± SEM. *P<0.05, **P<0.01, ***P<0.001, ****P<0.0001. P values calculated by nonparametric unpaired Mann-Whitney U test. Panels **A**, **D**, **L**, **N**, **and P** were created using BioRender.

To further evaluate the reconstituting potential of Kit^lo^ HSCs, we competitively transplanted Kit^lo^ or Kit^hi^ HSCs from young C57BL/6 (CD45.2) mice with rescue BM cells (CD45.1) into lethally irradiated young BALB/cJ recipients (**Figure 2D**). Eight weeks after allo-HCT, we found that Kit^lo^ HSCs generated significantly better B- and T-cell reconstitution, while myeloid reconstitution was comparable (**Figures 2E, S3A-S3B**). Concordant with a prior report of better self-renewal ability,^2^ we found that Kit^lo^ HSCs demonstrated increased LT HSC reconstitution (**Figure 2F**). In support of multilineage potential, we observed that Kit^lo^ HSC recipients had higher lymphoid progenitor reconstitution (including LMPP and CLP, **Figures 2F and 2G**), along with comparable myeloid precursor output (including MPP3, CMP, GMP, MEP, **Figures 2F and S3C**).

To evaluate whether enhanced lymphoid precursor output could contribute to thymic reconstitution, we concurrently evaluated the recipient thymi. We found that Kit^lo^ recipients demonstrated enhanced thymic recovery, characterized by significantly higher thymic cellularity and reconstitution of precursor and mature thymocytes (**Figures 2H-2J**). Notably, in support of a critical role for thymocyte-stromal crosstalk in preserving thymic architecture,^11, 12, 13^ we also found higher thymic epithelial cell (TEC), specifically medullary TEC, and endothelial cell recovery in Kit^lo^ reconstituted thymi (**Figures 2K and S3D**). We next examined donor-derived lymphoid progenitors for differential expression of molecules, involved in homing to the thymus (CCR7, CCR9, and PSGL1).^46^ We observed significantly higher expression of CCR7 on Kit^lo^-derived lymphoid progenitors (CLP) (**Figure S3E**). In addition, we analyzed thymic supernatants and found no differences in levels of thymopoietic ligands, IL22^47^ and RANKL^48^ (**Figure S3F**). Finally, we found significantly higher secondary lymphoid organ (spleen) reconstitution in Kit^lo^ recipients (**Figures S3G and S3H**). To evaluate long-term reconstitution, we performed additional harvests at 20-weeks following HCT. We observed similar trends in higher BM lymphopoiesis, thymic reconstitution, and thymic stromal recovery in Kit^lo^ recipients (**Figure S4**).

To assess thymic output, we used the RAG2-GFP model,^49^ which allows the analysis of recent thymic emigrants (RTEs) (**Figure 2L**). We observed higher numbers of PB GFP+ T cells in Kit^lo^ recipients **(Figure 2M**). Additionally, to assess whether this increased number PB T cells in Kit^lo^ recipients results in increased functionality, (**Figure 2N**), we analyzed response to ovalbumin engineered *Listeria Monocytogenes* (LM-OVA) and observed a significant increase in OT1+ CD8+ T cells (**Figure 2O**).

A key element in the post-HCT recovery of thymic function is the influx of bone marrow-derived lymphoid progenitors, also known as thymic seeding progenitors (TSPs).^10^ Furthermore, the significance of restoring the bone marrow-thymus axis in mitigating thymic involution has been underscored through various strategies aimed at augmenting BM-lymphopoiesis, including sex-steroid ablation^50, 51^ and administration of Ghrelin,^52^ among other approaches.^53^ We therefore hypothesized that robust BM lymphopoiesis driven by Kit^lo^ HSCs could potentially counteract age-related alterations within the thymic microenvironment, consequently ameliorating the decline in T-cell output. To this end, we competitively transplanted Kit^lo^ or Kit^hi^ HSCs from young C57BL/6 mice (CD45.2) with rescue BM cells (CD45.1) into lethally irradiated 14-month-old (Middle-aged) BALB/cJ recipients (**Figure 2P**). Sixteen weeks post-transplantation, we observed in Kit^lo^ vs Kit^hi^ recipients significantly higher numbers of peripheral T– and B-cells (**Figures 2Q, S3I, and S3J**). This was attributed to increased numbers of Kit^lo^ HSC-derived lymphoid progenitors (including LMPP/MPP4 and CLPs, **Figures S3K and S3L**), while numbers of myeloid precursors were comparable (MPP3, CMP, GMP, and MEP, **Figures S3K and S3M**). In our parallel analysis of the thymi of Kit^lo^ vs Kit^hi^ recipients, we observed increased numbers of thymocytes and TECs (**Figures 2R-2T and S3N**). Collectively, these findings establish the potential of Kit^lo^ HSCs for lymphopoiesis and to improve thymic recovery and peripheral T cell reconstitution in recipients of various ages.

### Kit^lo^ HSCs exhibit distinct epigenetic features that facilitate lymphoid potential

Prior work has shown that immunophenotypically defined HSC clones with differential lymphoid output have distinct epigenetic signatures, with no discernable patterns in terms of gene expression.^18, 24, 54^ To identify the gene regulatory network orchestrating the lymphoid potential differences in HSCs, we analyzed our single-cell ATAC-seq (scATAC-seq) component from our multiome dataset (**Figure S5D**). Through unsupervised clustering, we identified 6 clusters, including young and old HSCs (**Figures 3A and 3B**). Distinct HSC subsets identified via scRNA-seq analyses (**Figure 1B**) are situated in varied regions on the scATAC-seq UMAP (**Figures 3C and 3D**). Notably, MLin-HSC and Kit^lo^ HSC intersect with Cluster 3 from the scATAC-seq clusters (**Figure 3E**).

**Figure 3:**
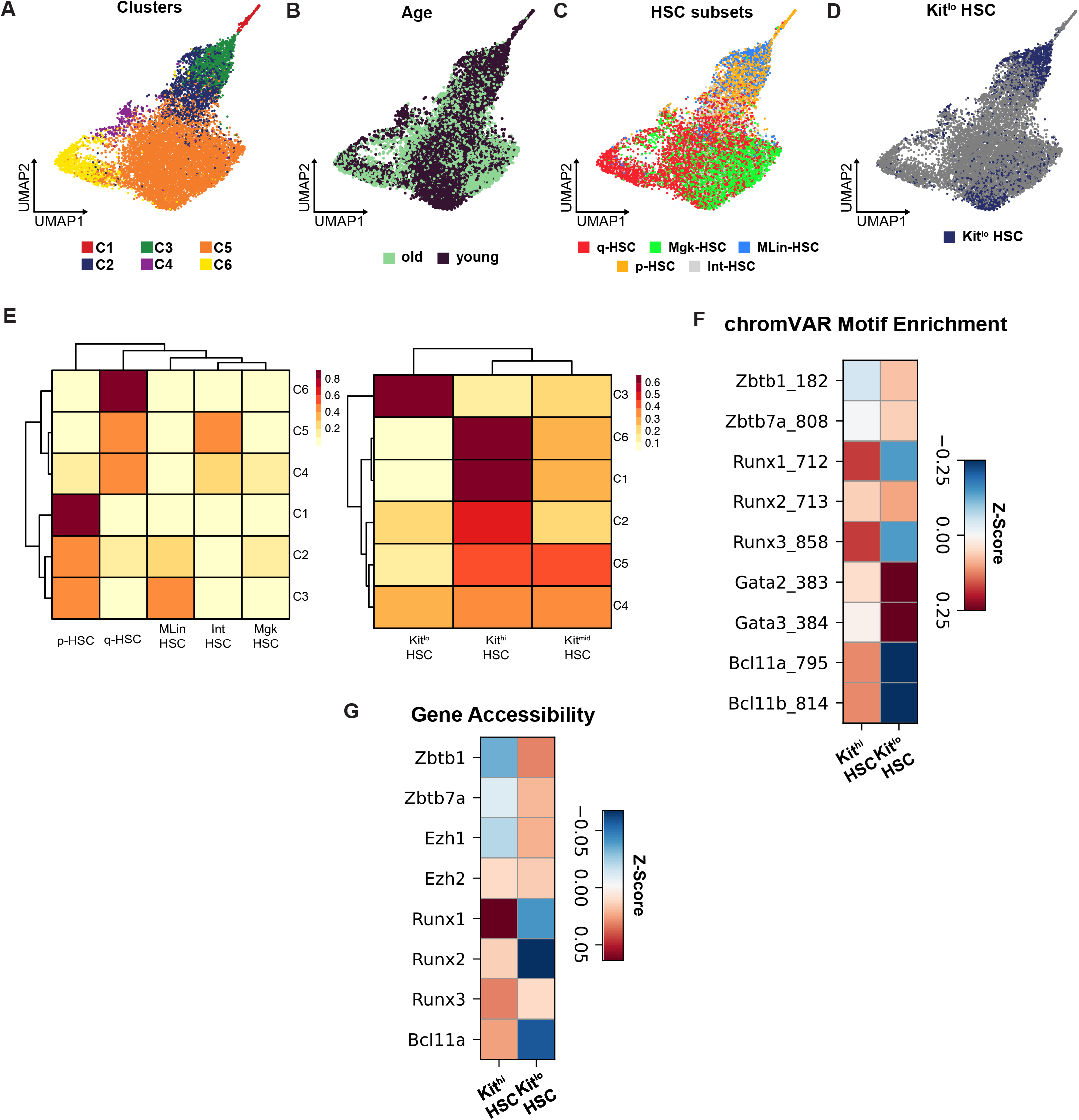
Kit^lo^ HSCs exhibit distinct epigenetic features that facilitate lymphoid potential. **(A)** UMAP of 6 unsupervised HSC clusters identified by tile-based scATAC-seq analysis after Harmony batch correction. **(B-D)** Annotated by age **(B)**, HSC subsets **(C)**, and Kit subsets **(D)** on the scATAC-seq UMAP. **(E)** scATAC-seq cluster composition by HSC subsets (**left**) and Kit subsets (**right**). (**F-G**) Matrix plots showing motif enrichment identified by ChromVAR (**F**) and gene accessibility (**G**) for lymphoid-specifying transcription factors for Kit subsets in young and old HSCs.

We characterized the differentially accessible regions (DARs) of chromatin that defined the HSC subsets and found 2816 DARs in cluster C3 (MLin-HSC and Kit^lo^ enriched) cluster compared to cluster C6 (q-HSC and Kit^hi^ enriched) (**Figure S5F**). Next to determine the differential activity and accessibility of lymphoid-specifying transcription factors (TFs) in the identified Kit^hi^ and Kit^lo^ HSC subsets, we analyzed chromatin accessibility variation within TF motifs using chromVAR and gene accessibility. Compared to Kit^hi^ HSCs, we found increased gene accessibility and enrichment for Zinc finger motifs, specifically *Zbtb1 and Zbtb7a,* in Kit^lo^ DARs, which are implicated in lymphoid differentiation program^18, 55, 56, 57^ (**Figures 3E-3F**).

In summary, we found that Kit^lo^ HSCs exhibit an epigenetic program, emphasizing an enrichment of lymphoid-specifying transcription factors, in support of their heightened lymphoid potential.

### *Zbtb1* regulates T-cell potential in phenotypic HSCs

Among members of the ZBTB gene family, *Zbtb1* plays a pivotal role in T-cell development^57, 58, 59^ and lymphoid reconstitution.^57^ However, its role in regulating lymphoid potential in HSCs remains unknown. We hypothesized that Zbtb1 loss impairs the lymphoid potential of Kit^lo^ HSCs. To this end, we used the CRISPR-Cas9 system to generate *Zbtb1*-deficient Kit^lo^ HSCs (**Figure 4A**) and confirmed decrease in Zbtb1 by flow-cytometric analysis (**Figure S6A**). *Zbtb1*-deficient Kit^lo^ HSCs showed significantly reduced *in vitro* T-cell potential (**Figure 4B**) and impaired lymphoid reconstitution *in vivo* (**Figure 4C**). Next, we questioned whether reinstating *Zbtb1* expression in megakaryocytic-biased, Kit^hi^ HSCs is sufficient to rescue their lymphoid defect. To test our hypothesis, we overexpressed *Zbtb1* cDNA in Kit^hi^ HSCs (**Figure 4D and S6B**) and we observed a rescue of their T-cell potential when compared to Kit^hi^ controls, thus partially phenocopying Kit^lo^ HSCs (**Figures 4E**) as well as improved lymphoid reconstitution *in vivo* (**Figure 4F**).

**Figure 4:**
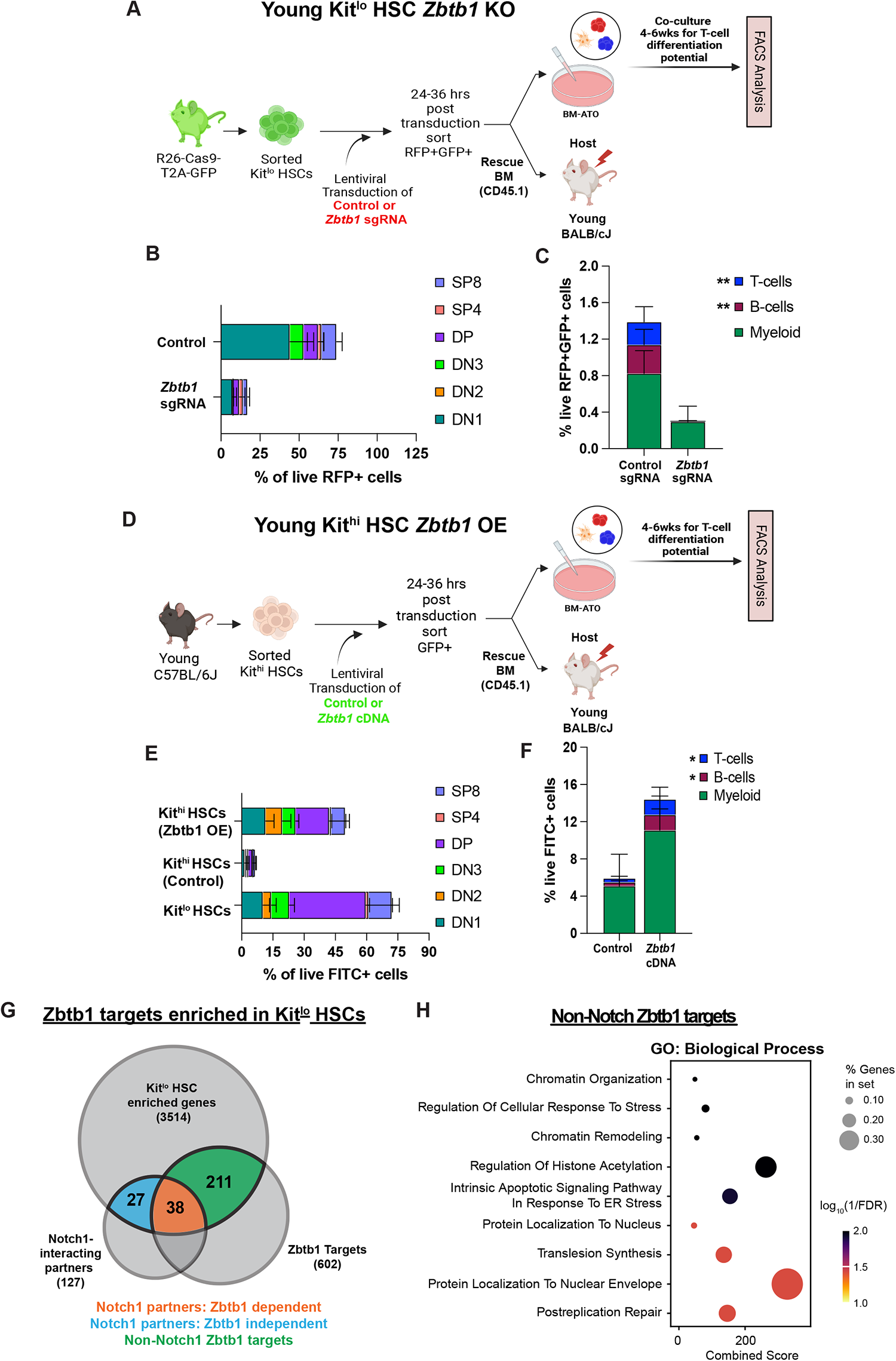
*Zbtb1* regulates T-cell potential in phenotypic HSCs. **(A)** Experimental schema for T-cell differentiation assay and competitive allogeneic HCT with *Zbtb1*-deficient Kit^lo^ HSCs generated with 2-mo young Rosa26Cas9-eGFP KI mice. **(B)** Frequency of T-cell subsets following 6-weeks of culture in mATOs. Aggregated data across 3 independent experiments. (C) Frequency of donor-derived lineage chimerism of control or *Zbtb1* KO Kit^lo^ HSCs in the PB at 6 weeks following competitive HCT. All data are from n=5-6 mice/group. (D) Experimental schema for T-cell differentiation assay and competitive allogeneic HCT with *Zbtb1*-OE Kit^hi^ HSCs from young mice. (E) Frequency of T-cell subsets following 6-weeks of culture in mATO of young Kit^hi^ HSCs. Aggregated data across 3 independent experiments. (F) Frequency of donor-derived lineage chimerism of control or *Zbtb1* OE young Kit^hi^ HSCs in the PB at 6-weeks following competitive HCT. All data are from n=5-6 mice/group. (G) Previously identified Zbtb1-targets^60^ and Notch1-interacting proteins^62^ enriched in Kit^lo^ HSCs. (H) Pathway enrichment analysis for non-Notch1 Zbtb1-targets enriched in Kit^lo^ HSCs by *gseapy*()^91^ using GO_Biological_Process_2023 libraries. Bubbleplot showing representative pathways. Error bars represent mean ± SEM. *P<0.05, **P<0.01, ***P<0.001. P values calculated by nonparametric unpaired Mann-Whitney U test. Panels **A** and **D** were created using BioRender.

Previous studies have identified Zbtb1 targets which facilitate Notch-mediated T-cell differentiation in lymphoid progenitors^60^ and support chromatin remodeling following DNA damage.^61^ Given *Zbtb1* motif enrichment and enhanced T-cell differentiation in Kit^lo^ HSCs, we hypothesized that Zbtb1 targets mediating lymphoid differentiation gene programs are also enriched in Kit^lo^ HSCs. To this end, we queried Kit^lo^ enriched genes for Notch1-interacting proteins.^62^ We observed ∼50% overlap of Notch1-partners enriched in Kit^lo^ HSCs, of which 60% were Zbtb1-targets identified in a previous CHIP-seq study^60^ (**Figure 4G**). Furthermore, we identified an additional subset of Zbtb1 targets (33%) that were not associated with the Notch1 interactome (**Data Table 3**). To further query for non-Notch related biological pathways, we performed pathway analysis and identified a strong enrichment for gene sets related to chromatin remodeling and RNA stability (**Figure 4H, Data Table 4**).

These data demonstrate that Zbtb1 contributes to an epigenetic regulatory program that facilitates T-cell differentiation in primitive Kit^lo^ HSCs likely through instructive and permissive mechanisms.

### Old Kit^lo^ HSCs exhibit enhanced T-cell potential

To determine whether Kit^lo^ HSCs retain their lymphoid potential with age, we evaluated the lymphoid potential of old Kit subsets. Remarkably, even in the case of old mice, we noted a substantial increase in lymphoid progenitor and ATO output from old Kit^lo^ HSCs compared to Kit^hi^ HSCs (**Figures 5A-5B**), although modest when compared to the young HSCs. This finding is consistent with prior reports of proliferative defects in aged T-cell precursors contributing to decreased thymopoiesis. ^3, 4, 5^

**Figure 5:**
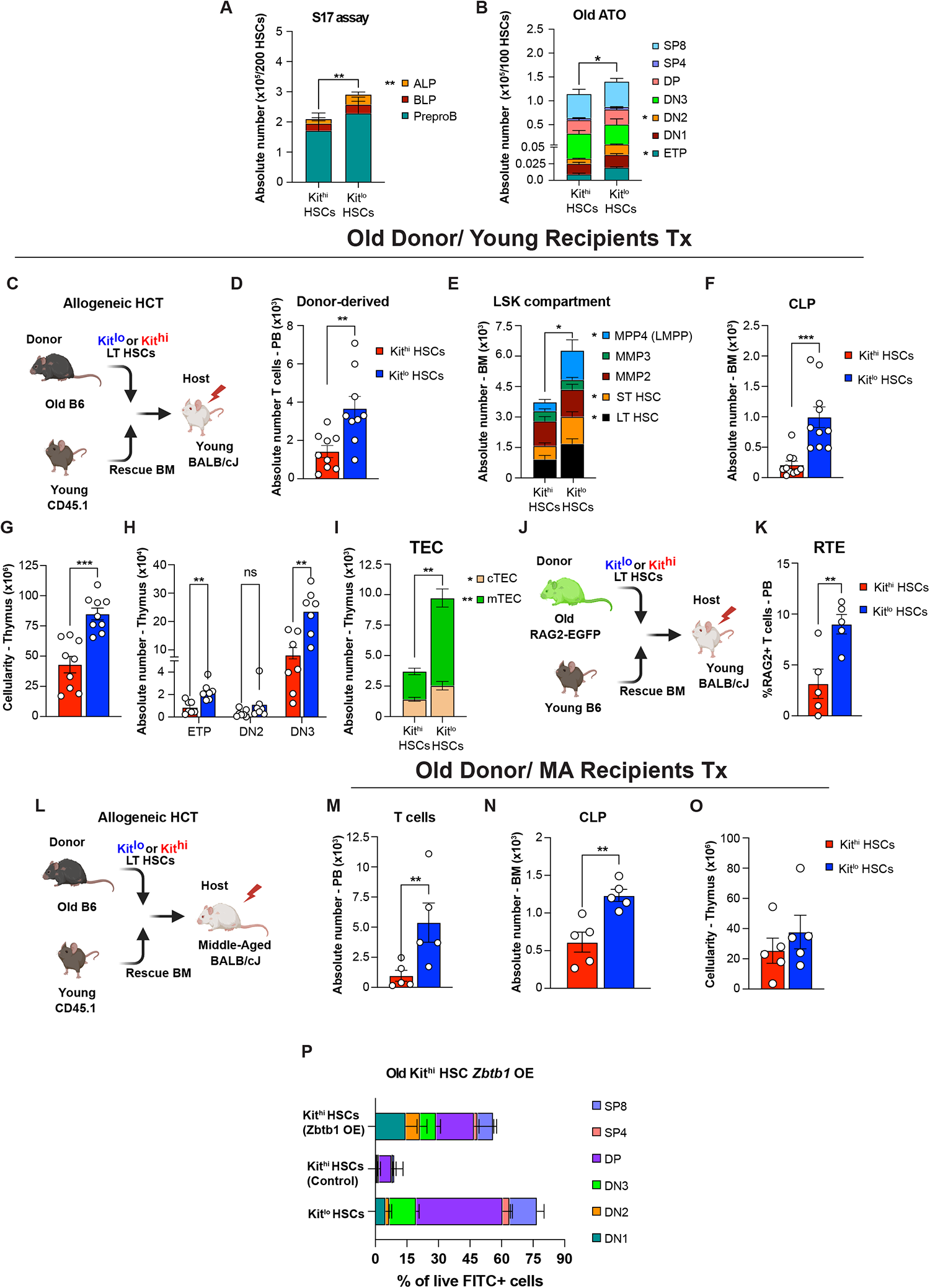
Old Kit^lo^ HSCs exhibit enhanced T-cell potential. **(A-B)** Evaluation of *in vitro* lymphoid progenitor and T-cell differentiation potential of Kit^hi^ (red) and Kit^lo^ HSCs (blue) from or 24-mo (old) C57BL/6 mice using the S17 lymphoid assay and murine artificial thymic organoid (mATO) respectively, as shown in Figure 2A. Enumeration of absolute number of lymphoid progenitor cells following 14 days in S17 lymphoid assay **(A)** and absolute number of T-cell subsets following 8-weeks of culture in mATOs **(B).** Aggregated data across 4-5 independent experiments. (**C)** Experimental schema for competitive allogeneic HCT (allo-HCT) using Kit^hi^ (red) and Kit^lo^ HSCs (blue) from 22–24-mo (old) C57BL/6 mice with competitor bone marrow (BM) cells from B6.SJL-PtprcaPepcb/BoyJ mice transplanted into lethally irradiated young BALB/cJ recipients. **(D-I)** Eight weeks after competitive HCT enumeration of absolute number of donor-derived T-cells in the PB **(D)**, LSK cell subsets as defined in Figure 3C (**E)**, CLP cells as defined in Figure 3D **(F)**, total thymic cellularity **(G)**, donor-derived T-cell precursor thymocyte subsets as defined in Figure 3F **(H)**, TEC compartment as defined in Figure 3H **(I).** All data are from n=9-10 mice/group. (**J)** Experimental schema to evaluate thymic function following competitive allo-HCT using old HSCs. **(K)** Frequency of donor-derived Recent Thymic Emigrants (RTE: CD3^+^GFP^+^) in the PB at 8 weeks post-HCT. All data are from n=5 mice/group. (**L)** Experimental schema for competitive allogeneic HCT (allo-HCT) using Kit^hi^ (red) and Kit^lo^ HSCs (blue) from 22-24-mo (old) C57BL/6 mice with competitor bone marrow (BM) cells from B6.SJL-PtprcaPepcb/BoyJ mice transplanted into lethally irradiated middle-aged BALB/cJ recipients. **(M-O)** Eight weeks after competitive HCT enumeration of absolute number of donor-derived T cells in PB **(M)**, donor-derived CLPs in the BM as defined in Figure 3D **(N)**, and thymic cellularity **(O)**. All data are from n=5-6 mice/group. (**P**) Frequency of T-cell subsets following 6-weeks of culture in mATO of old Kit^hi^ HSCs following *Zbtb1* OE. Aggregated data across 3 independent experiments. Error bars represent mean ± SEM. *P<0.05, **P<0.01, ***P<0.001, ****P<0.0001. P values calculated by nonparametric unpaired Mann-Whitney U test. Panels **C, J,** and **L** were created using BioRender.

Next, to test their *in vivo* potential, we competitively transplanted either Kit^lo^ or Kit^hi^ HSCs from old C57BL/6 (CD45.2) mice with rescue BM (CD45.1) cells into lethally irradiated young BALB/cJ recipients (**Figure 5C**). At eight weeks post-transplantation, we noted significantly higher numbers of T cells in the peripheral blood of Kit^lo^ vs Kit^hi^ recipients. (**Figures 5D and S7A**). Consistent with our experiments using HSCs from young donors, we observed in the BM of Kit^lo^ vs Kit^hi^ recipients significantly higher numbers of LT HSCs, LMPPs and CLPs (**Figures 5E-5F**) with comparable myeloid potential (**Figures 5E and S7B**). Furthermore, mirroring our findings in HSCs from young mice, we observed in recipients of old Kit^lo^ HSCs enhanced overall thymic cellularity, thymocyte reconstitution, and recovery of TECs and endothelial cells (**Figures 5G-5I and S7C-S7D**), increased CD4+ and CD8+ T cells in the spleen (**Figures S7E and S7F**), and higher RTE output (**Figures 5J-5K**). At 20-weeks post-HCT, we observed similar trends of enhanced reconstitution in old Kit^lo^ HSC recipients, confirming their long-term reconstituting potential (**Figure S7H-S7O**). Next to examine whether old HSC subsets with preserved lymphoid could augment aged thymic recovery, we competitively transplanted middle-aged recipients with old Kit^lo^ or Kit^hi^ donors (**Figure 5L**). We observed increased CLPs in the BM and peripheral T cells, but a modest, non-significant increase in thymic cellularity in Kit^lo^ recipients (**Figures 5M-5O, and S7G**). Similar to our *Zbtb1* functional studies with young Kit^hi^ HSCs, we observed a rescue in T-cell potential in *Zbtb1*-overexpressed old Kit^hi^ HSCs (**Figure 5P**).

Finally, to directly compare the per cell functionality of Kit^lo^ HSCs in the context of aging, we generated mixed chimeras combining Kit^lo^ HSCs cells from both young and old mice in an equal competition transplant (**Figure S8A**). Consistent with our *in vitro* findings (**Figures 5A-5B**), we found an overall decrease in lymphoid reconstitution within the PB, BM, and thymus with old Kit^lo^ HSCs when compared to their young counterparts (**Figures S8B-S8G**). Pathway analysis revealed decreased enrichment of chromatin remodeling related gene sets (**Figure S8H, Data Table 5**). Additionally, we noted decreased accessibility and expression of *Zbtb1 (***Figures S8I-S8K**) in Old Kit^lo^ HSCs, indicating a impaired epigenetic program for lymphoid differentiation. These findings demonstrate that old Kit^lo^ HSCs exhibit preserved lymphoid potential and contribute to thymic recovery, albeit with reduced output.

### KIT^lo^ human BM HSCs exhibit enhanced lymphoid potential

Informed by a recent study that mapped the molecular profiles of human thymic seeding progenitors (TSPs) to their transcriptional counterparts within the HSC compartment,^63^ we next sought to identify a comparable human HSC subset with lymphoid potential. Interrogating a CITE-Seq dataset^64^ generated from young and old human bone marrow (BM) samples (**Figures 6A-6C, S9A**), we mapped our mouse Kit^lo^ gene signature onto human HSCs and noted a significant enrichment in young BM (**Figures 6D and 6E**). We observed lower protein expression of KIT (CD117) on young HSCs by using antibody-derived tag (ADT) reads (**Figure S9B**). To validate our in-silico findings, we next assessed KIT expression in human BM samples (young: 20-40 years; middle-age/ old: >40 years). In our FACS analysis, we found that phenotypic HSCs (pHSCs: CD34+CD38-CD10-CD45RA-CD90+) showed the lowest KIT expression (**Figure S9C**), with higher KIT levels on old pHSCs (**Figure 6F**). Moreover, we noted a statistically significant association between age and the composition of KIT subsets in pHSCs, indicating an increase in KIT^hi^ and a decrease in KIT^lo^ HSCs, aligning with our observations in mice. (**Figure 6G**).

**Figure 6.**
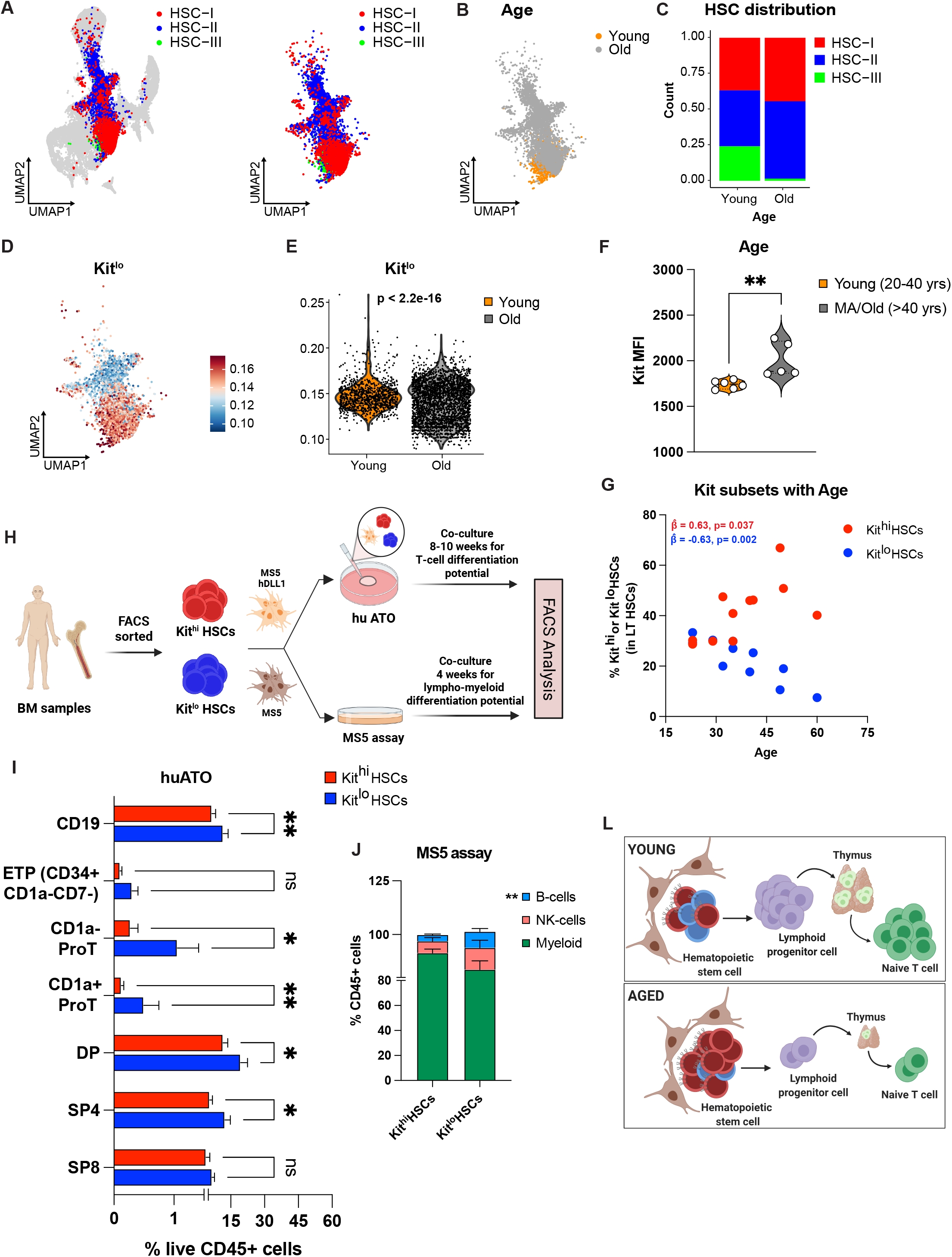
KIT^lo^ human BM HSCs exhibit enhanced lymphoid potential. **(A-E)** Young and old human BM CITE-seq dataset generated and published by Sommarin et al.^64^ **(A-C)** UMAP of CD34+ young and old human BM annotated with HSC subsets **(A)**, and age **(B)**. **(C)** HSC cluster composition by age. **(D)** Kit^lo^ gene signature (top 50 marker genes from our mouse data converted to human orthologues) was overlaid on human HSC UMAP. **(E)** Violin plot for Kit^lo^ gene score by age. Statistical analysis was performed using the Wilcoxon test. **(F-G)** FACS analysis of human BM samples. **(F)** Violin plot of KIT protein expression on phenotypic HSCs (pHSC: CD34+CD38-CD10-CD45RA-CD90+) by age (young: 20-40yrs; MA/old: >40yrs). **(G)** Scatterplot showing distribution of KIT^hi^ and KIT^lo^ HSC subsets within pHSCs in human BM samples with varying age. **(H)** Experimental schema to evaluate *in vitro* T-cell and lympho-myeloid potential of human BM HSC subsets based on differential KIT expression using human artificial thymic organoids (hATO) and MS5 assay respectively. **(I)** Following 8-10-weeks of culture, enumeration of CD19+ B-cells and T-cell subsets within CD34+ T-cell precursors (Early Thymic Progenitors, ETP: CD34^+^CD1a^-^CD7^-^; CD1a^neg^-ProT: CD34^+^CD1a^-^CD7^+^; CD1a^pos^-ProT: CD34^+^CD1a^+^CD7^+^), and mature T-cells (DP: CD34^-^ CD5^+^CD7^+^CD4^+^CD8^+^; SP4: CD34^-^CD5^+^CD7^+^CD4^+^CD8^-^; SP8: CD34^-^CD5^+^CD7^+^CD4^-^CD8^+^). Aggregated data represent mean from 8 independent BM donors ± SEM. **(J)** Following 4-weeks of culture, enumeration of Lineage output (Neutrophil-Granulocytes: hCD45+CD15+; monocyte-macrophages: hCD45+CD14+; B-cells: hCD45+CD19+; NK-cells: hCD45+CD56+) after culturing 200 KIT^lo^ and KIT^hi^ HSC cells on MS-5 stroma, as previously described^67^. Aggregated data represent mean from 5 independent BM donors ± SEM. **(L)** Model illustrating age-related loss of lymphoid potential is attributed to a decreased in the frequency of Kit^lo^ HSCs in aged BM. Error bars represent mean ± SEM. *P<0.05, **P<0.01, ***P<0.001, ****P<0.0001. P values for panel **G** was based on generalized estimating equation logistic model adjusted for age, KIT^lo^ and KIT^hi^ groups and their interaction, the matched data structure captured by an exchangeable working correlation structure. P values calculated either by nonparametric paired Mann-Whitney U test (Panels **I** and **J**), unpaired Mann-Whitney U test (Panel **F**). Panels **H** and **L** were created using BioRender.

Cord blood (CB) and BM have previously been fractionated based on KIT levels.^65, 66^ To evaluate the T-cell potential of these human BM HSC subsets, we generated artificial human thymic organoids (huATOs) using purified KIT^hi^ or KIT^lo^ HSCs from human BM samples (**Figures S9D and 6H**; **Data Table 6**). In concordance with our mouse *in vitro* studies, following 8-10 weeks of culture, we observed significantly higher output of precursor and mature T-cells from KIT^lo^ HSCs-derived ATOs compared to KIT^hi^ HSCs (**Figure 6I**). To further evaluate the multilineage potential of KIT HSC subsets, we performed a lympho-myeloid differentiation assay, as previously described.^67^ While we observed comparable myeloid differentiation potential, Kit^lo^ HSCs exhibited significantly more B cell and a non-significant increase in NK cell differentiation potential (**Figure 6J**).

Collectively, these results identify a previously uncharacterized subset of human BM HSCs, KIT^lo^ HSCs – which exhibit enhanced lymphoid potential.

## DISCUSSION

Recent studies have uncovered significant heterogeneity within the HSC compartment and their role in hematopoiesis. This heterogeneity has implications for disease development and treatment. Of note, certain aspects of HSC heterogeneity challenge the established hematopoietic models, with lineage biases and lineage-restricted cells impacting self-renewal dynamics.^68, 69^ Decisions regarding lympho-myeloid fate are established at multiple levels of the hematopoietic hierarchy: HSCs,^21, 22, 70^ MPP, LMPP, and more mature GMPs and CLPs. At the level of HSCs, the process of epigenetic imprinting confers a distinct advantage for lymphoid priming within multilineage HSCs, as well as maintaining these cell autonomous features imposed at steady state, even under conditions of stress.^18, 24^

Here, we report on the enhanced T-cell potential of Kit^lo^ HSCs, observed in both mice and humans. Through combining multiomic single-cell sequencing and functional analyses utilizing a preclinical allo-HCT model, we demonstrate augmented potential of Kit^lo^ HSCs for T-cell lymphopoiesis and to support thymic recovery, independent of age. During the early post-transplant period, we noted a relative increase in DN3 thymocytes relative to other thymocyte precursor subsets. This observation is consistent with previous studies reporting that DN3 thymocytes undergo compensatory expansion to support early post-transplant thymopoiesis, with minimal contribution from T-cell precursor thymocytes (ETP and DN2).^10, 71, 72^ Further investigation into mechanisms driving these compensatory processes and its implications for immune function is critical to our understanding of post-transplant immune reconstitution.

Mechanistically, Kit^lo^ HSCs exhibit differential expression and activity of lymphoid-specifying transcription factors (TFs), including *Zbtb1*, indicating their epigenetic program towards a lymphoid fate. Studies in mouse models have shown that disruptions in *Zbtb1* result in T-cell developmental defects and severe combined immunodeficiency phenotypes,^57, 58^ along with competitive disadvantages in B- and NK-cell compartments. ^57^ In our study, we identify *Zbtb1* as an essential transcription factor governing lymphoid specification in phenotypic HSCs and facilitating T-cell differentiation by reprogramming lineage biased Kit^hi^ HSCs. Additionally, we observed functional differences within Kit^lo^ HSCs in the context of aging, specifically a competitive disadvantage in Old Kit^lo^ HSCs, potentially attributable to their differences in expression of the epigenetic repressor of multipotency, *Ezh1,*^73^ and *Zbtb1*. Consistent with prior research, our study suggests *Zbtb1* modulates Notch-dependent^60^ and cellular survival programs.^29, 74^ However, further mechanistic studies are needed to elucidate the precise transcriptional mechanisms underlying this regulation. While the definitive role of *Zbtb1* in HSCs is not established, it may not have a direct role at the HSC stage; instead, its expression could be indicative of epigenetic states that facilitate or permit Zbtb1 expression and function at later stages of development, in lymphocyte progenitors. Future work will also address if ZBTB1 could be used to augment generation of mature lymphoid lineages from human BM precursors.

Long-term reconstituting CD34+KIT^lo^ human BM cells have been previously reported.^66^ In contrast, fractionating phenotypic HSCs in Cord Blood (CB) based on KIT levels showed reduced reconstitution of KIT^int^ HSCs compared to KIT^hi^ HSCs.^65^ These findings suggest that differences in Kit signaling have functional relevance depending on the developmental stage. In our study, combining transcriptional, immunophenotypic, and functional analyses of human BM samples we identified a KIT^lo^ subset within immunophenotypically defined HSCs with enhanced lymphoid potential, highlighting its translational relevance in human biology. Further functional characterization is warranted to definitively establish the clinical significance of thse observations. This understanding is crucial as prospective HSC isolation with subsequent ex vivo expansion,^75, 76^ holds promise for enhancing immune regeneration after bone marrow transplantation, effectively counteracting treatment-related immunosuppression and age-associated thymic decline.

## ACKNOWLEDGEMENTS

We thank Dr. Andrea Schietinger for the gift of mice; Ronan Chaligné, Kalina Belcheva, and Anthony Michaels for help; Dr. Gay Crooks, and Julia Gerscheimer (UCLA) for sharing the mATO and huATO systems and assistance with troubleshooting; Kenneth Dorshkind (UCLA) for providing S17 cells; Parashar Dhopala and Göran Karlsson (Lund University) for providing the human BM CITE-seq raw matrices.^64^

## AUTHOR CONTRIBUTIONS

HKE, MRMvdB conceived and designed the study. HKE, MRMvdB acquired funding. HKE, MBdS, AR, BG, NL, MAEA, SG planned research activities. HKE curated data and performed initial analysis. AIK and SM performed initial computational analysis of single-cell RNA-seq data, and single-cell ATAC-seq and human CITE-seq data, respectively. RS provided code and assisted HKE with downstream data/sequence computation and statistical analyses. HKE analyzed flow cytometric data. HKE, MBdS, AR, BG, NL, MAEA, XZ, HA, SG, MS, SDW, KV, CYP, CED, and JCS planned and performed experiments. CSL, AB, MRMvdB supervised study. HKE, SM, and MBdS prepared figures and visualized data. HKE, MRMvdB wrote the initial draft. All authors reviewed and approved the manuscript.

---

**Figure S1.**
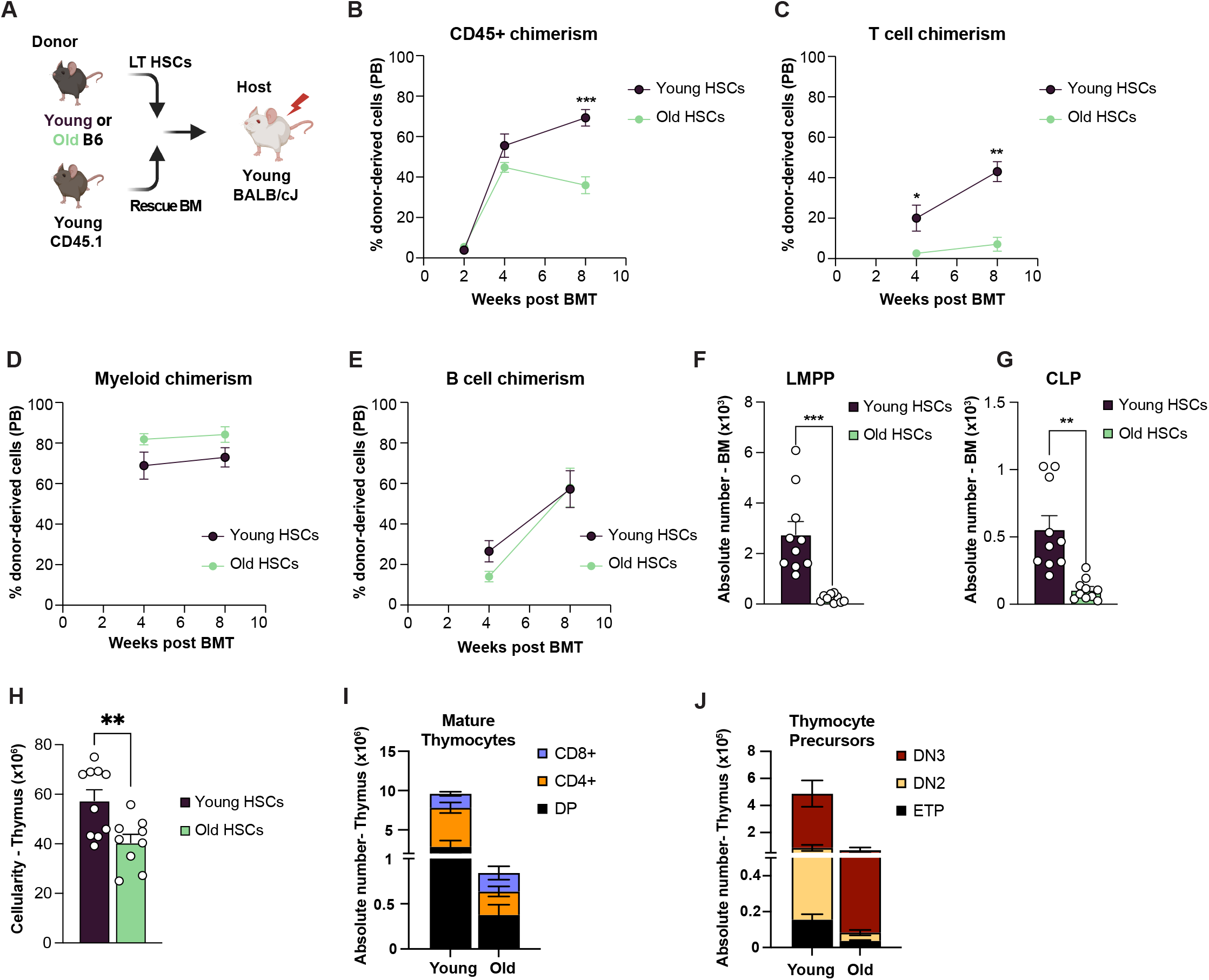

**Figure S2.**
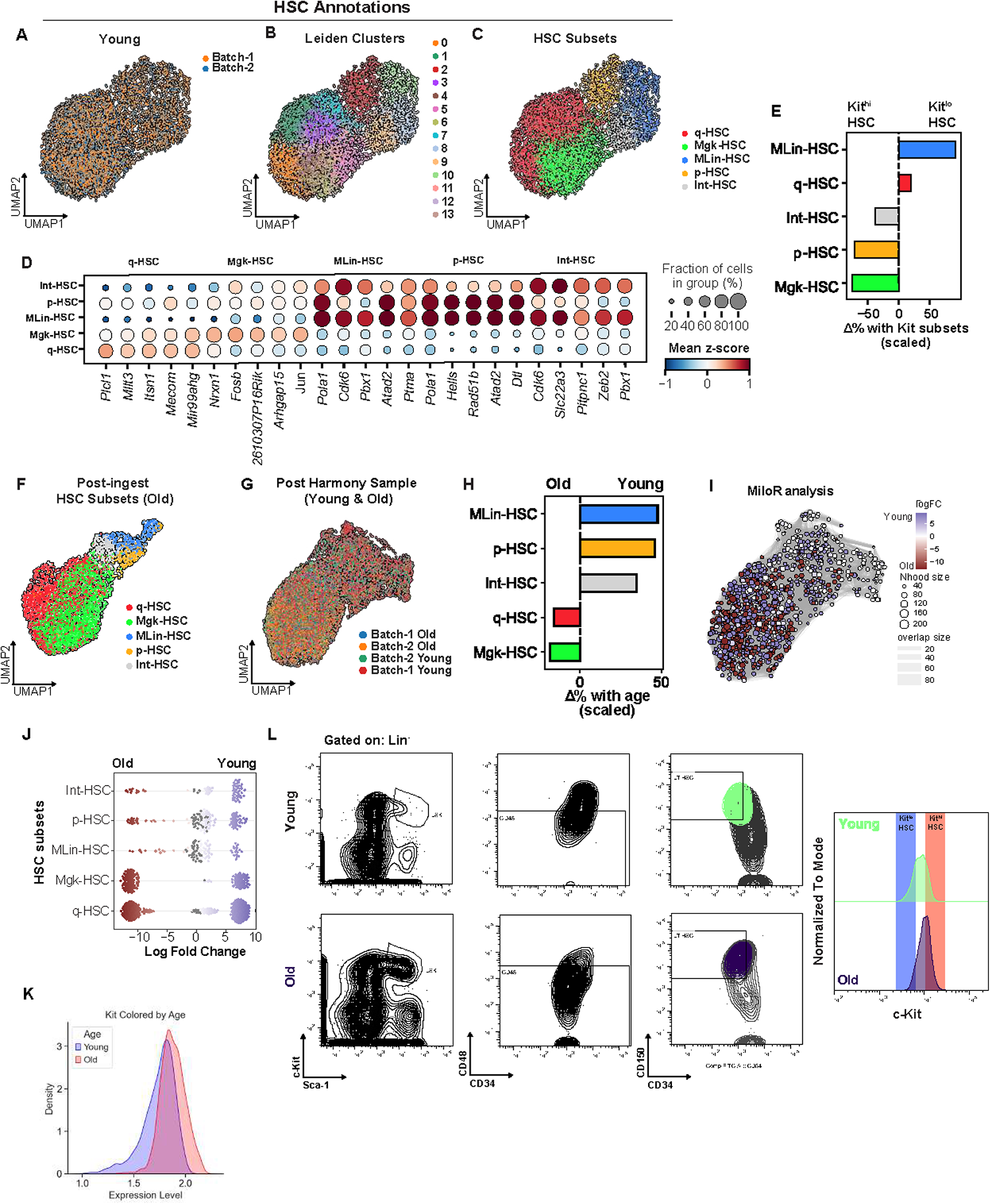

**Figure S3.**
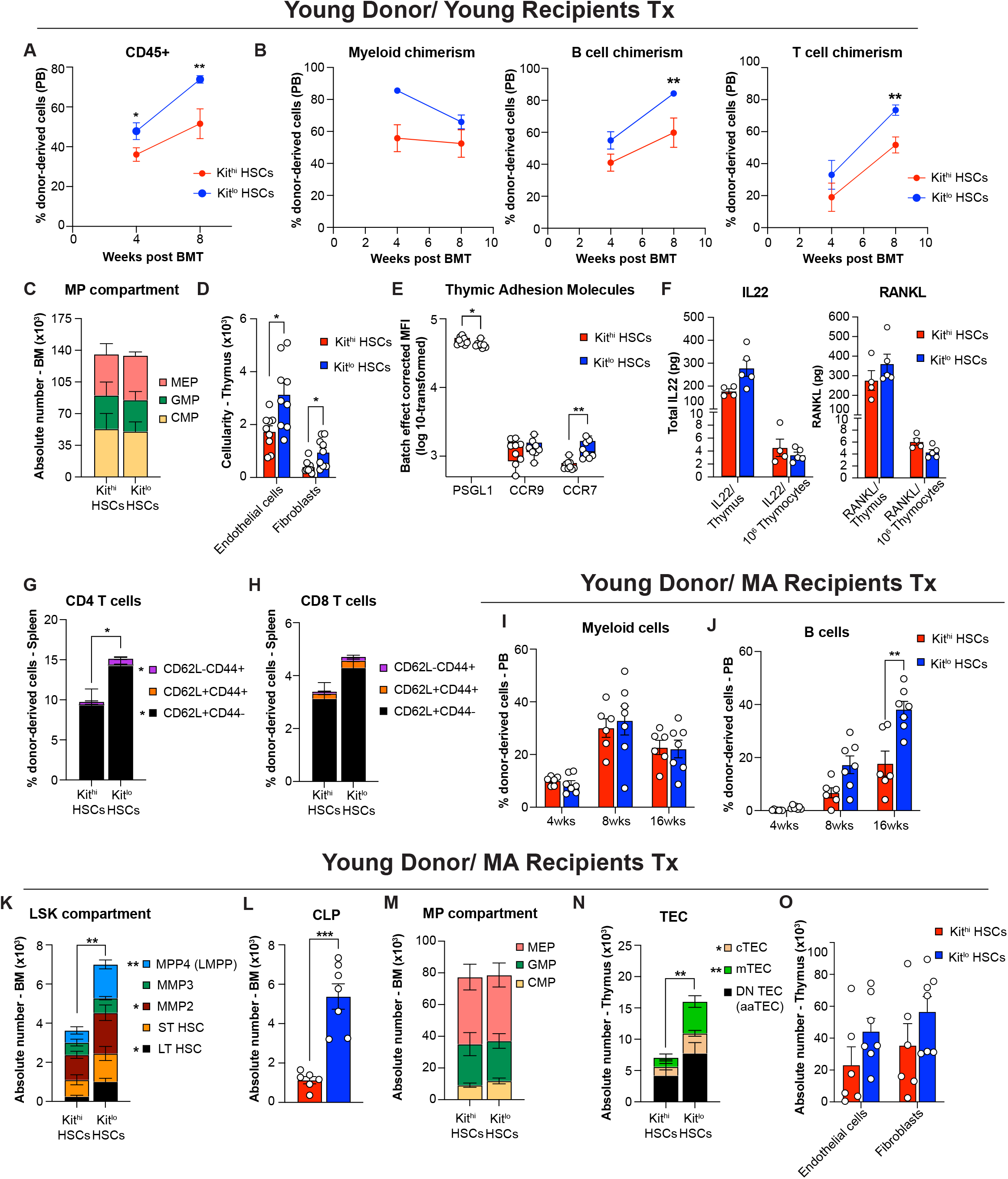

**Figure S4.**
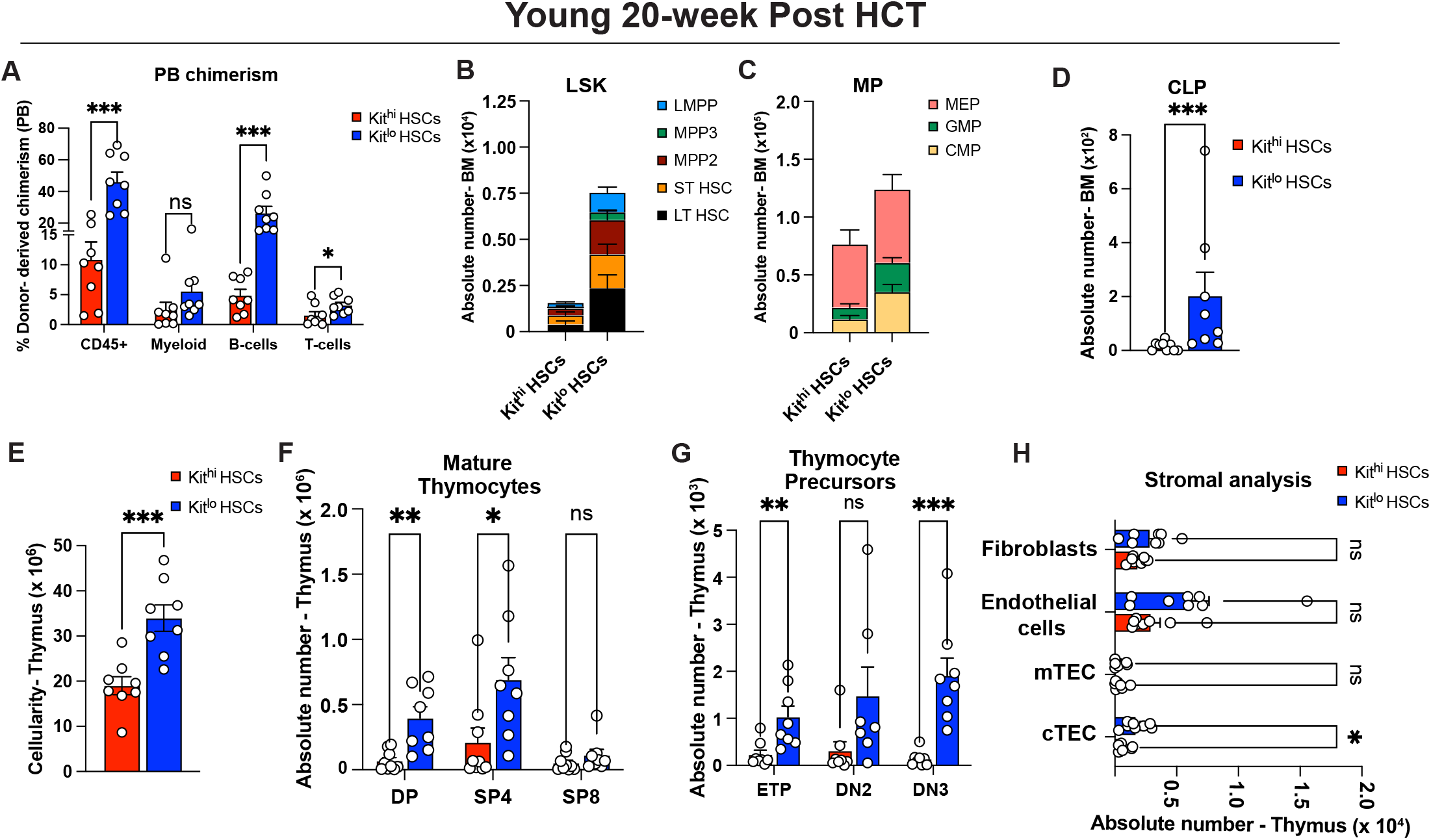

**Figure S5.**
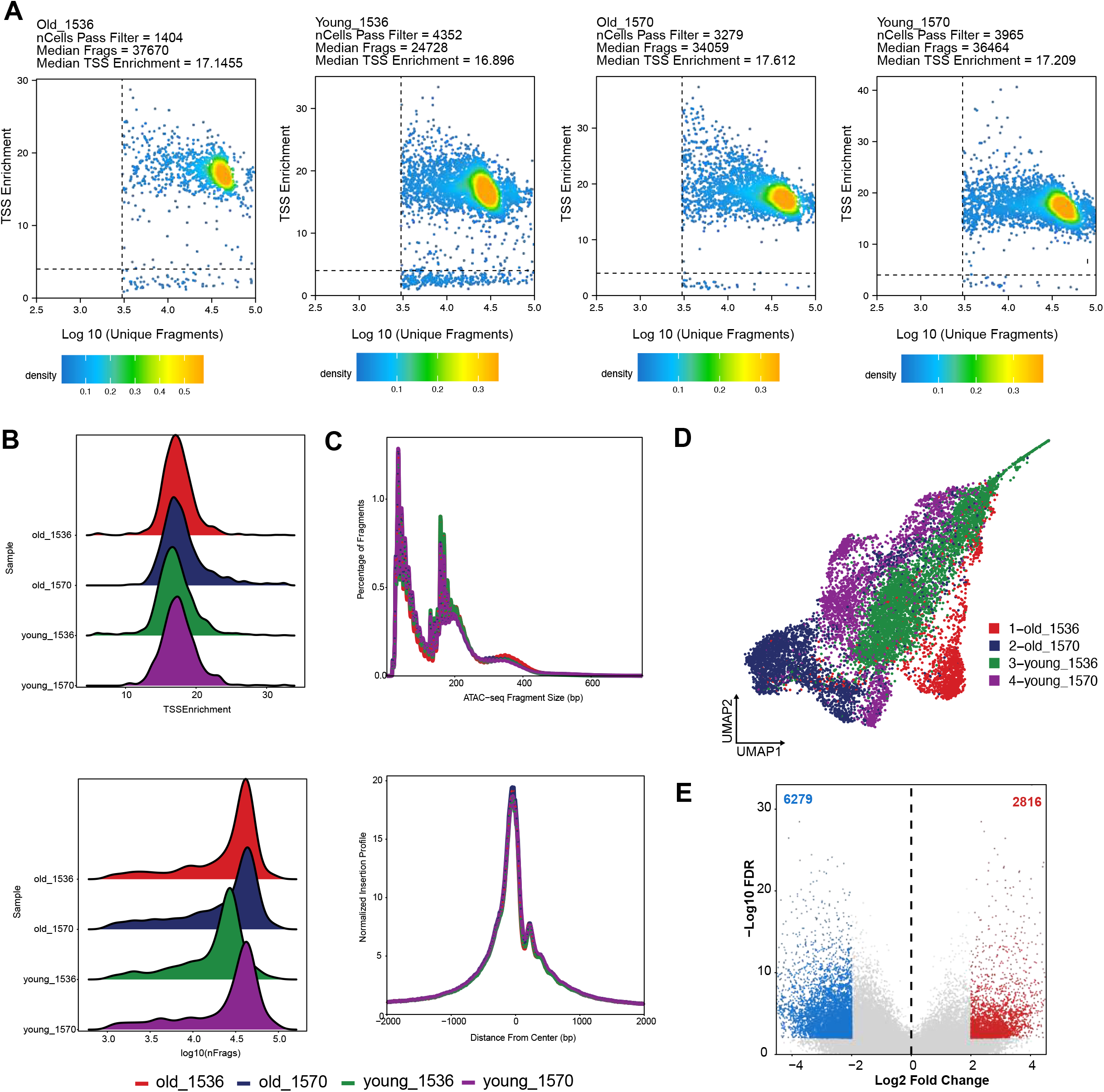

**Figure S6.**
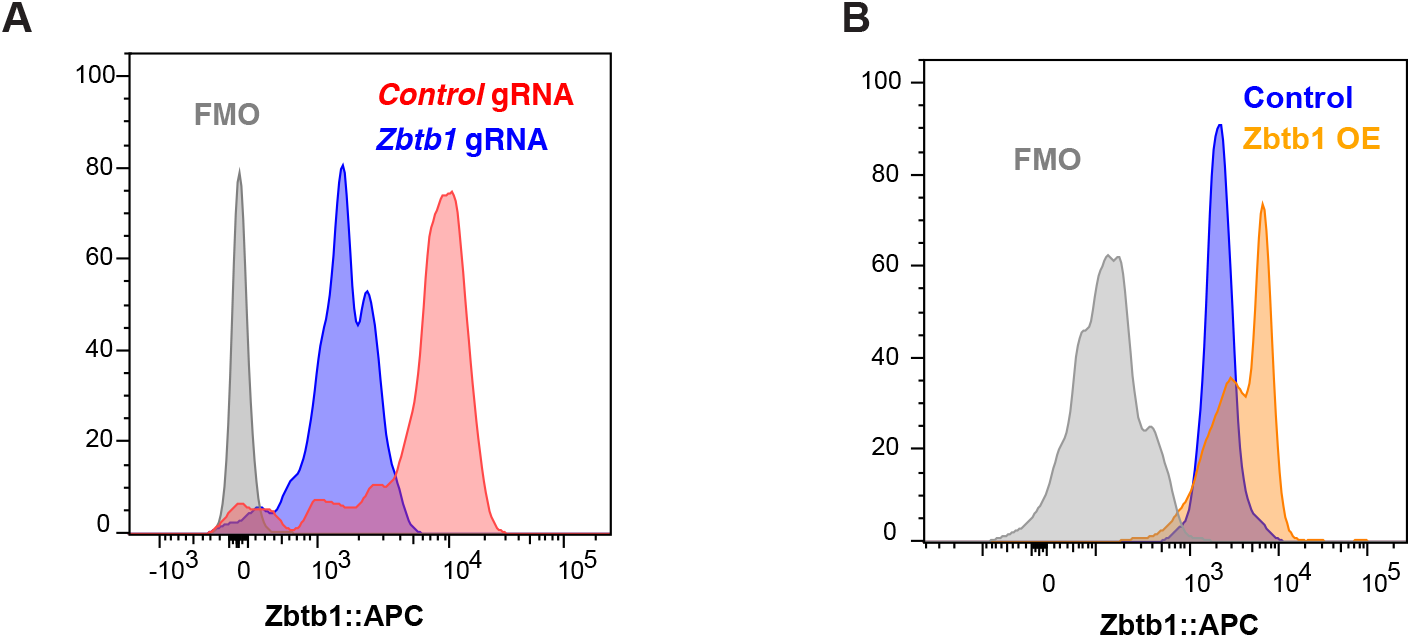

**Figure S7.**
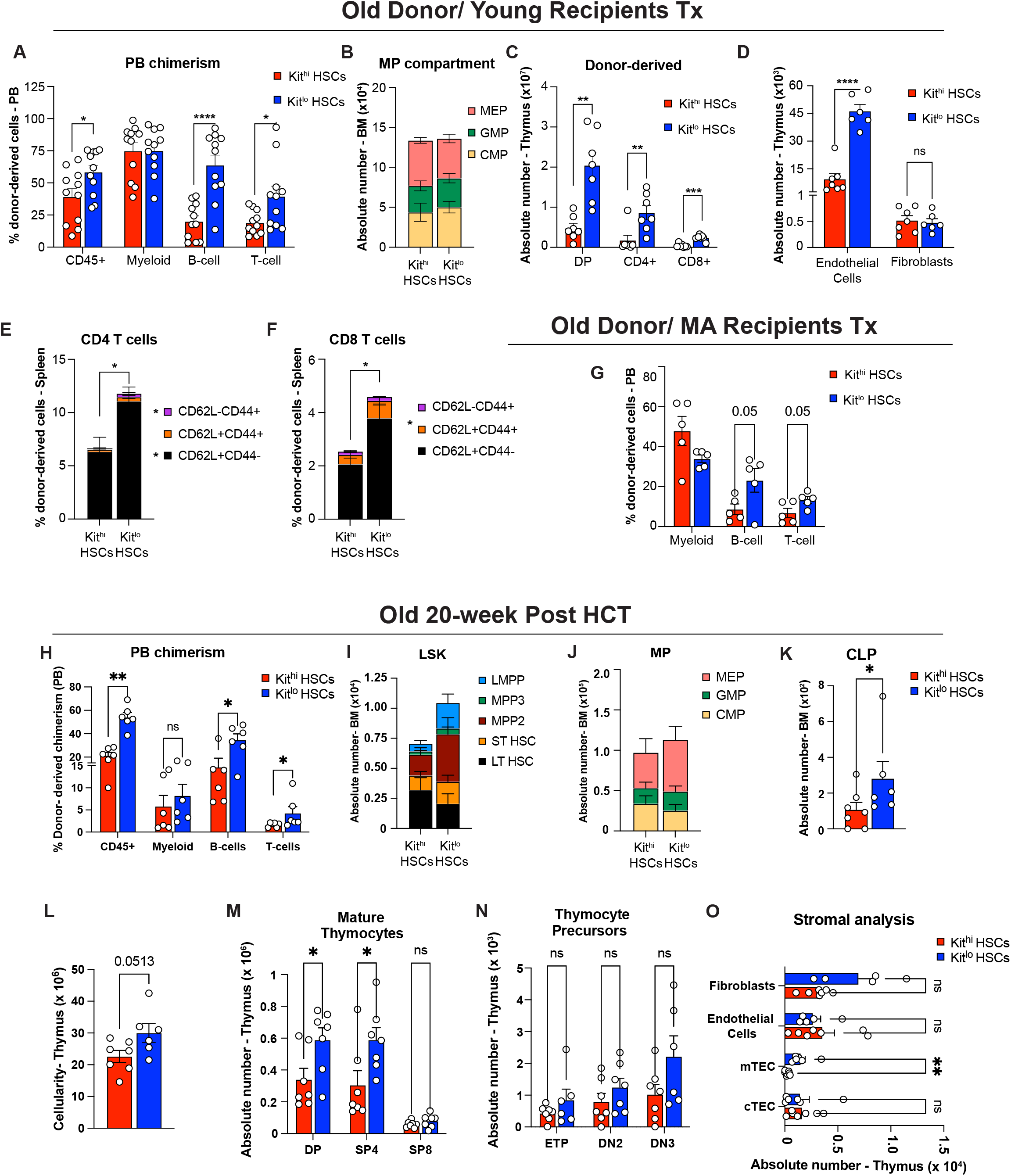

**Figure S8.**
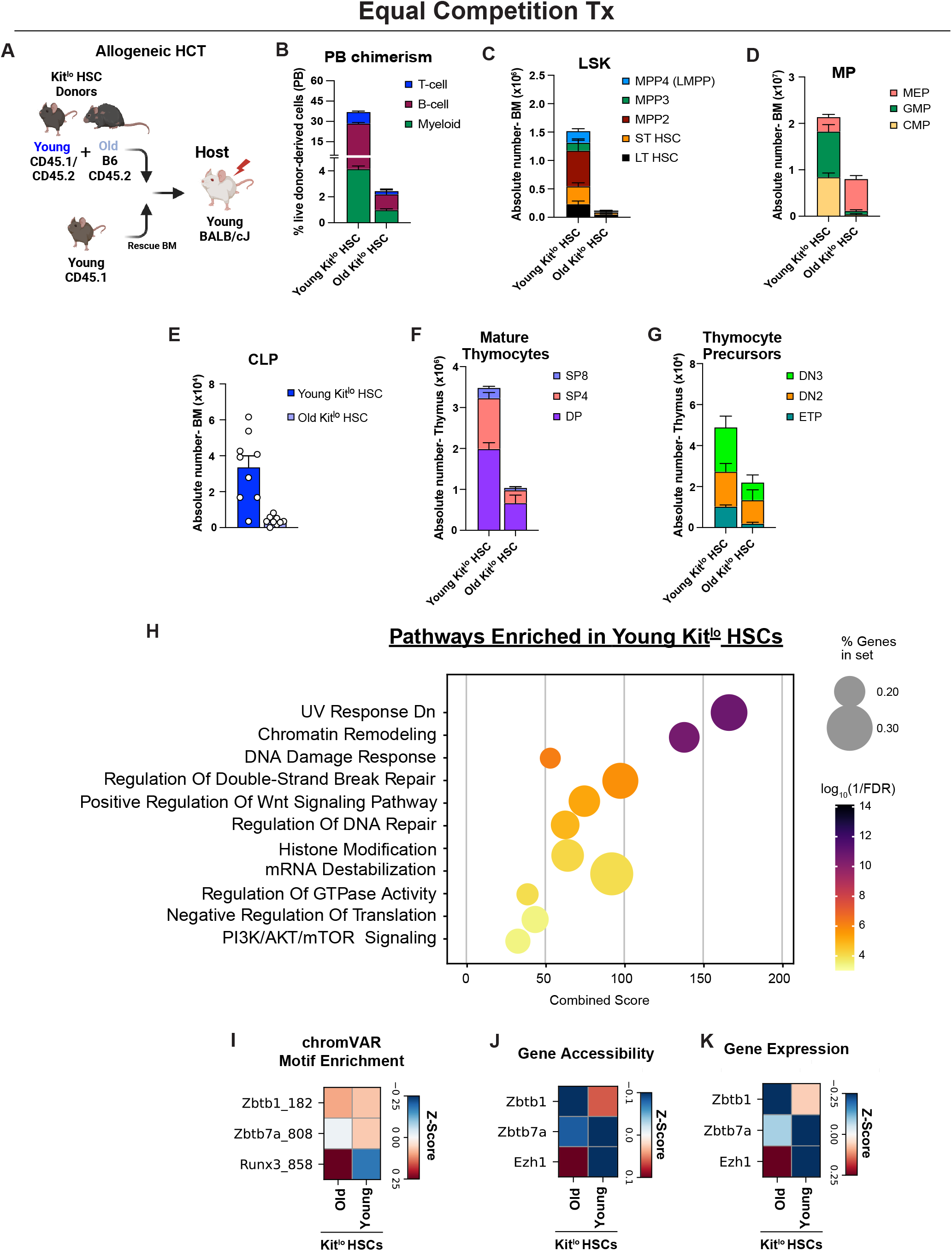

**Figure S9.**
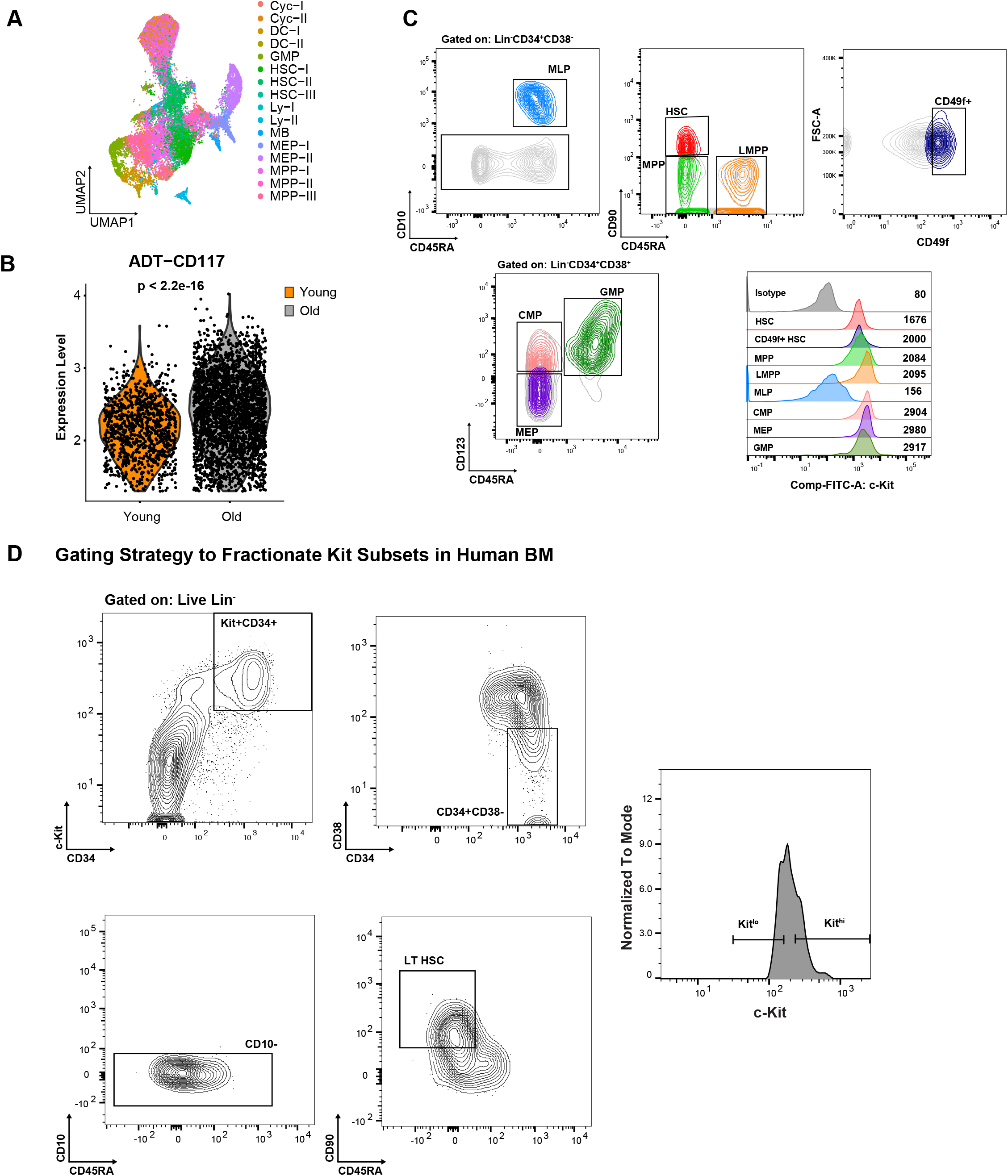

