## Supplementary material for "An epigenetically distinct HSC subset supports thymic reconstitution": Methods and Supplemental Figure legends

**Animal use**

Young mice (6-10 weeks of age): C57BL/6J (CD45.2/H-2Kb), B6.SJL-PtprcaPepcb/BoyJ (CD45.1), BALB/cJ (H-2Kd), and B6(C)-Gt(ROSA)26Sor^em1.1(CAG-cas9*,-EGFP)Rsky/J^ (Rosa26^Cas9^ KI ) mice were purchased from The Jackson Laboratory. Aged C57BL/6 (CD45.2/H-2Kb) mice (23-24 months of age) were obtained from National Institute of Aging (Baltimore, MD). Aged mice (ranging between 18-24 months of age) correlate with humans ranging between 56-69 years of age. We obtained middle-aged mice (ranging between 14-16 months of age) by initially purchasing young BALB/cJ (H-2Kd) mice from Jackson Laboratories (JAX) and subsequently allowing them to age under controlled conditions in our facility. Middle-aged mice (ranging between 14-16 months of age) correlate with humans ranging between 40-60 years of age. RAG2-EGFP-CD45.1 chimeric mice were generated by crossing FVB-Tg (RAG2-EGFP)1Mnz/J (JAX# 005688) and B6.SJL-PtprcaPepcb/BoyJ (CD45.1, JAX# 002014). Young OT-1-CD45.2 (Va2+Vb2+/ CD45.2/H2-Kb) were obtained from Dr. Andrea Schietinger. For consistency all experiments were carried out using only female mice. These mice were allowed to acclimatize in our vivarium for at least 10-14 days before experiments. Mice were euthanized with CO2 gas inhalation and were maintained under pathogen-free conditions according to an MSKCC IACUC-approved protocol.

**Cell lines**

To provide a microenvironment that supports lymphoid progenitor in vitro, we cocultured purified HSCs from young and old B6 bone marrow with the S17 stromal cells^1^, which were originally obtained from K. Dorshkind (UCLA). To support in vitro T-cell differentiation, we co-cultured

purified HSCs from young and old B6 bone marrow and human BM with MS5-mDLL4 or MS5-hDLL4 stromal cells ^2^, which were obtained from Dr. Gay Crooks (UCLA). Subsequently, both cell lines were maintained in our laboratory as described in the original references. Details of the differentiation cultures are given below under HSC Lymphoid Differentiation Assays.

**Cell Preparation and Staining**

Femurs, tibias, pelvises, and the spine were dissected from euthanized mice and cleaned of muscles and connective tissue on ice in 1X PBS. Bone marrow cells were collected by crushing all the bones with a sterile mortar and pestle in FACS buffer: 1X PBS containing 2% Fetal Bovine Serum (Hyclone) and 2 mM EDTA. BM cells were enriched for c-Kit+ cells using mouse CD117 magnetic microbeads (Miltenyi Biotec) per manufacturer’s protocol.

**Bone Marrow analysis and Flow-cytometric Isolation of Hematopoietic Stem Cells**

BM cells were stained with fluorochrome-conjugated antibodies as well as propidium iodide (Molecular Probes) or Zombie Aqua™ Fixable Viability Kit (Biolegend). All monoclonal antibodies were purchased from Biolegend and eBiosciences. The monoclonal antibodies included Phycoerythrin-Cyanine 7 (PE-Cy7) antibodies to lineage (Lin) markers (CD3ε, CD4, CD8a, B220, Gr-1, Mac-1 and Ter119), Brilliant Violet 711 (BV711) or Brilliant Violet 605 to Sca-1, APC/Cyanine7 to cKit, FITC antibody to CD34, PE antibody to CD150 (SLAM), Pacific Blue to CD48, APC to CD45.2 and PerCP/Cyanine5.5 to CD45.1. Cells were analyzed or sorted using a FACS LSR II UV or Aria II cell sorter (BD Biosciences). For all experiments (transplantation and in vitro assays) in which Kit^hi^, Kit^lo^ or Kit HSCs were purified, they were double sorted to ensure >95% purity.

To calculate bone marrow cellularity and frequency of donor-derived hematopoietic precursors, two femurs and two tibias of primary recipient (BALB/cJ) were flushed into FACS buffer at 8- or 20-weeks post-transplant. Collected cells were then incubated with red blood cell lysis buffer (ACK lysis buffer, Thermo Fisher Scientific) for 8 minutes and then washed twice with PBS/2.5% FBS. Cells were resuspended and then stained in PBS/2.5% fetal calf serum with fluorochrome-conjugated antibodies (Biolegend, BD, eBiosciences) against lineage (Lin) markers (CD3ε, CD4, CD8a, B220, Gr-1, Mac-1 and Ter119) (PE-Cy5), Sca-1 (BV711), c-Kit (APC-Cyanine7), CD150/ SLAM (BV605), CD135/Flt3 (BV421 or PE-Cy5), CD127/IL7Ra (APC), CD48 (BV510), CD16/32 (AF-700), CD34 (FITC), CD45.2 (BUV395), CD45.1 (PE-Cy7 or AF-700), H-2Kd (PE) and H-2Kb (BV650 or BV786). For additional thymic adhesion molecule analyses, antibodies against CCR7 (BV421), CCR9 (FITC) and PSGL1 (PE) were used for chimerism studies. Following a 30-45 incubation on ice, cells were washed in in PBS/2.5% fetal calf serum. Finally, propidium iodide (Molecular Probes) was added as a viability dye before acquiring the data. Cells were analyzed or sorted using a FACS LSR II UV (BD Biosciences).

**Transplantation experiments**

**A. For competitive transplantation assays**, double-FACS sorted Kit^hi^, Kit^lo^ HSCs cells from young or old CD45.2 mice were mixed with unfractionated BMMCs from young CD45.1 mice. Cells were transplanted via retro-orbital sinus injections into lethally irradiated young (6-8-week-old) or middle aged (14-16-month-old) BALB/cJ recipient mice (9 Gy, single dose, using a 79 X-Ray source) under isoflurane anesthesia, within 1h post-irradiation. **B. For Kit^lo^ equal competition experiment,** 375 double-FACS sorted Kit^lo^ HSCs cells from both young (CD45.1/CD45.2) and old CD45.2 mice were combined with 5x10^5^ unfractionated BMMCs from young CD45.1 mice and transplanted into lethally irradiated 6-8-week-old BALB/cJ recipient mice. **C. For RTE experiments**, 750 double-FACS sorted Kit^hi^, Kit^lo^ HSCs cells from young and old RAG2-GFP (CD45.1) mice were mixed with 5x10^5^ unfractionated BMMCs from young CD45.2 mice and transplanted into lethally irradiated 6-8-week-old BALB/cJ recipient mice. **C. OT-1 Transplant and Adoptive T-cell transfer for functional assessment**, 750 Kit^hi^ and Kit^lo^ HSCs were double FACS-sorted from 8–10-week-old OT-1 mice and competitively transplanted with 5x10^5^ unfractionated competitor BMMCs from (CD45.1) into lethally irradiated young (6-8-week-old) BALB/cJ (H-2Kd) recipients. Eight-week post-transplant (BMT), spleens of primary recipients were brought into single-cell suspension and 1/10 of splenocytes were adoptively transferred into young (8-10wk) C57BL/6 secondary recipients. 4 hours later, mice were infected with 5.000 CFU L. monocytogenes bacteria expressing chicken ovalbumin (LM-OVA) to evaluate functional CD8+T cell response to primary infection. On day 8 post-infection, spleens of infected mice were analyzed via flow cytometry. Counting beads were used to calculate absolute cell numbers.

**Peripheral blood analysis**

Peripheral blood samples were collected in 50 mM EDTA solution (Thermo Fisher Scientific) via retro-orbital sinus bleeds. Thereafter PB was incubated with red blood cell lysis buffer (ACK lysis buffer, Thermo Fisher Scientific) for 8 minutes and then washed twice with PBS/2.5% fetal bovine serum. Cells were resuspended and then stained in PBS/2.5% fetal calf serum with antibodies against CD3 (AF700), B220 (PE-Cy7), Gr-1 (APC), Mac-1 (APC), NK1-1 (BV605), CD45.2 (BUV395), CD45.1 (BV421), H-2Kd (PE) and H-2Kb (FITC) for chimerism studies for 30 mins followed by wash in in PBS/2.5% FBS. Finally, propidium iodide (Molecular Probes) was added as a viability dye before acquiring the data. Cells were analyzed using a FACS LSR II UV and LSR Fortessa X50 (BD Biosciences). To generate absolute T-cell counts, complete blood counts, including differentials, were obtained using a ProCyte Dx Hematology Analyzer (IDEXX).

**Thymi Harvests: FACS and ELISA analysis**

All steps were performed on ice unless indicated. Primary recipient (BALB/cJ) thymi at 8 or 20 wks post-transplant were excised and enzymatically digested following an adapted protocol.^3^ Briefly, thymi were mechanically dissociated into ca. 2 mm pieces. Tissue pieces were incubated with a digestion buffer (RPMI, 10% FCS, 62.5 um/mL liberase TM, 0.4 mg/ml DNase I) twice for 30 min at 37 C. Between incubation steps, supernatant containing dissociated cells was transferred to 50 mL conical tubes equipped with 100 um filter.

For thymic immunophenotypic analysis, cells were first incubated with Fc block solution (anti-CD16/CD32 antibody) for 10 min on ice. Solution was discarded and cells were stained in PBS/2.5% fetal calf serum with fluorochrome-conjugated antibodies against lineage (Lin) markers (CD19, CD11b, NK1-1, TCRγδ, Gr-1, Ter119) (Biotin), Streptavidin (PE-TexasRed), CD4 (PE-Cy7), CD8 (BV711), CD25 (BV510), CD44 (AF700), c-Kit (APC), CD135/Flt3 (BV421), CD45.2 (BUV395), CD45.1 (BV605), H-2Kd (PE) and H-2Kb (FITC). for chimerism studies. To characterize the CD45- compartment, thymic cells were stained with antibodies against UEA-1 (FITC), 6C3 (PE), EPCAM (BV605), PDGFRa (BV421), MHC-II (APC), CD31 (PE-Cy7), CD45 (BUV395) and Ter-119 (PE-Cy5.5) for 30 mins followed by wash in in PBS/2.5% FBS. Antibodies were purchased from either (Biolegend and eBiosciences), except Ulex europaeus agglutinin 1 (UEA-1), conjugated to FITC, was purchased from Vector Laboratories (Burlingame, CA). Finally, 7-AAD (Molecular Probes) was added as a viability dye before acquiring the data. Flow Cytometric analysis was performed on FACS LSR II UV (BD Biosciences).

For all ELISA experiments, thymi were suspended by mechanical dissociation. The resultant supernatants were quantified using mouse cytokine (IL-22 and RANKL) specific ELISA kits from R&D Systems and read on a Spark Multimode Microplate Reader (TECAN).

**Spleen Harvest and Analysis**

All steps were performed on ice unless indicated. Primary recipient (BALB/cJ) spleens at 8- or 20-weeks post-transplant were excised and crushed, filtered through a 70uM strainer (pluriSelect), intermittently washed with PBS/2.5% fetal bovine serum and collected in a 50ml conical tube. Subsequently, spleen pellet was incubated in red blood cell lysis buffer (ACK lysis buffer, Thermo Fisher Scientific) for 8 minutes and then washed twice with PBS/2.5% fetal bovine serum. Prior to staining, an aliquot was set aside for counts with the Nexcelom Cellometer K2. For spleen immunophenotypic analysis, cells were first incubated with Fc block solution (anti-CD16/CD32 antibody) for 10 min on ice. Solution was discarded and stained in PBS/2.5% fetal calf serum with fluorochrome-conjugated antibodies against lineage (Lin) markers (CD19, CD11b, NK1-1, TCRγδ, Gr-1, Ter119) (Biotin), Streptavidin (PE-TexasRed), CD4 (PE-Cy7), CD8 (BV711), CD62L (BV510), CD44 (AF700), CD45.2 (BUV395), CD45.1 (BV605), H-2Kd (PE) and H-2Kb (FITC). for chimerism studies.

**HSC Lymphoid Differentiation Assays**

**A. S17 Progenitor Assay**

Lymphoid progenitor production from young and old B6 bone marrow was measured by mixing FACS purified 200 Kit^hi^ and Kit^lo^ HSCs with 50,000 S17 stromal cells^1^ in 1.5 mL of methylcellulose (MC) medium. MC medium was prepared by supplementing a-MEM with 30% heat-inactivated FCS, 1% methylcellulose (STEMCELL Technologies), 5x10-5 M 2ME, 2 mM L-glutamine, 50 µg/mL gentamicin, 100 U/mL streptomycin, 100 µg/mL penicillin, 0.1 mM MEM vitamins, 0.1 mM nonessential amino acids, 1 mM sodium pyruvate, 20 ng/mL stem cell factor, 20 ng/mL Flt3L ligand, and 50 ng/mL IL-7 (all from Biosource). The mixture was plated in non-tissue-culture treated 3.5-cm2 dishes (Becton Dickinson). Following 12 days of culture, the contents of the plates were harvested, cells were enumerated, and examined for production of lymphoid progenitors by flow cytometry. All cultures were placed at 37C, 5% CO2 humidified incubators until processing.

**B. Mouse and Human ATO Assay**

Prior to setting up the ATO experiments, the MS5 stromal cell lines were thawed and expanded in culture for at least 7 days. The cell lines and ATOs were set up and maintained as initially described.^2^

For the mouse ATO system, FACS purified 100 Kit^hi^ and Kit^lo^ HSC cells from young and old mice were combined with 150k MS5-mDll4 stromal cells to form each murine ATO. The media (DMEM/F-12, 50x B27, 100x Glutamax, 100x Pen/Strep, and 1000x Ascorbic acid) was changed twice a week, with beta-mercaptoethanol (1000x) and cytokines (5ng/mL murine IL-7, 5ng/mL Flt3L, and 5ng/mL SCF) added fresh each time.

For the human ATO system, healthy human bone marrow mononuclear cells were purchased from BIOIVT (Johnson City, TN) for FACS sorting of Kit^hi^ and Kit^lo^ HSCs within phenotypic HSCs (CD34^+^CD38^-^CD10^-^CD45RA^-^CD90^+^). The sorted cells were then aggregated with MS5-hDll4 (50-150 HSCs and 150k MS5-hDll4 cells per ATO) and seeded on culture inserts (Millipore Sigma). The human ATO media (RPMI 1640, 25x B27, 100x Glutamax, 100x Pen/Strep, 1000x Ascorbic Acid) was refreshed twice a week, supplemented with 2.5ng/mL recombinant human IL-7, 5ng/mL recombinant human Flt3L (Peprotech), and 5ng of human SCF was used only for the first week of differentiation.

For analysis via FACS, the ATO was mechanically disrupted, filtered through a 70uM strainer (pluriSelect), and counted with the Nexcelom Cellometer K2 prior to staining. Harvest timepoints varied, dependent on the initial primary cell population (mATO: 6-8-weeks; huATO: 8-10-weeks).

**C. Lympho-myeloid Differentiation Assay**

As previously described^4^, MS-5 cells were seeded onto 0.1% gelatin-coated 24-well plate at an initiating density of 2 × 10^4^/well in α-MEM medium (Thermo Fisher) supplemented with 10% FBS (Gibco), 1% penicillin-streptomycin (Hyclone) and 1% GlutaMAXTM (Gibco). Twenty-four hours after plating of MS-5 stroma, 200 cells were added into each well in the presence of 0.1 μM DuP-697 (Cayman Chemicals), 20 ng/ml SCF (Peprotech), 10 ng/ml G-CSF (Peprotech), 10 ng/ml FLT3L (Peprotech), 10 ng/ml IL-2 (Peprotech), 10 ng/ml IL-15 (Sino Biological). Cultures were maintained for 4 weeks with weekly half media changes (with 2x cytokines). Cocultures were transferred onto fresh MS-5 stroma every two weeks through 40 μm filter to remove the stromal cells. All the cells in each well were harvested and analyzed by flow cytometry at week 4. The antibodies used to read the lineage outputs in this assay were anti-human CD45-BV480, anti-human CD14-BUV805, anti-human CD15-BV421, anti-human CD19-FITC, anti-human CD56-BUV496, and anti-human CD235a-PE-Cy7.

**Lentiviral Production**

For deletion experiments, scramble and Zbtb1 guide RNAs (sgRNA#1: GTTTTAGAGCTAGAAATAGCAAGTTAAAATAAGGCTAGTCCGTTATCAACTTGAAAAAGTGGCACCGAGTCGGTGC; sgRNA#2: GTTTTAGAGCTAGAAATAGCAAGTTAAAATAAGGCTAGTCCGTTATCAACTTGAAAAAGTGGCACCGAGTCGGTGC) were cloned into U6-EFS-mCherry (VectorBuilder), while Control and mouse Zbtb1 [NM_178744.3] were individually cloned into CMV-EFS-EGFP (VectorBuilder) for over-expression experiments and utilized for lentivirus generation. Lentivirus was generated by cotransfecting the above plasmids containing either sgRNAs or cDNA, pMD2.G (Addgene #12259), and psPAX2 (Addgene#12260) into HEK 293 T cells. SFEM Media was replaced 6h post-transfection, and the virus suspension was harvested 36-48 h post-media change. The supernatant was collected and used for lentiviral transductions.

**HSC Lentiviral Transduction**

As previously outlined sorted Kit^lo^ and Kit^hi^ HSCs from young, aged B6, and young Rosa26^Cas9^ KI mice were cultured overnight in SFEM (STEMCELL Technologies) media supplemented with murine cytokines (50 ng/ml SCF, 10 ng/ml TPO) in single-wells of a 96-well plate (TC treated). The cells were transduced with lentiviral suspensions by spinfection in retronectin (Takara Bio)-coated plates for 90 mins. Post spinfection, cytokine-supplemented fresh media was added to the transduced wells. Twenty-four hours later, cells were either sorted for RFP and GFP positivity (for deletion experiments), or GFP expression (for over-expression experiments) using the BD Aria instrument. KO and OE was confirmed by flow cytometric analysis with antibodies against Zbtb1 (AlexaFluor647, Signalway Antibody Cat#C47476-AF647).

**Single Cell Multiome ATAC and Gene Expression Cell Preparation**

Single Cell Multiome ATAC + Gene Expression was performed with the 10X genomics system using Chromium Next GEM Single Cell Multiome Reagent Kit A (catalog no. 1000282) and ATAC Kit A (catalog no. 1000280) following Chromium Next GEM Single Cell Multiome ATAC + Gene Expression Reagent Kits User Guide and demonstrated protocol - Nuclei Isolation for Single Cell Multiome ATAC + Gene Expression Sequencing. Briefly, cells (viability 95%) were lysed for 4min and resuspended in Diluted Nuclei Buffer (10x Genomics, PN- 2000207). Lysis efficiency and nuclei concentration was evaluated on Countess II automatic cell counter by trypan blue staining. Nuclei were loaded per transposition reaction, with a targeting recovery between 1,000 and 10,000 nuclei after encapsulation. After transposition reaction nuclei were encapsulated and barcoded. Next-generation sequencing libraries were constructed following User Guide, which were sequenced on an Illumina NovaSeq 6000 system.

**SEQUENCING DATA PROCESSING**

**Single Cell RNA-seq**

**i) Preprocessing and downstream data analysis**

FASTQ files were processed using the 10x Cell Ranger package (v7.01). The Cell Ranger generated filtered_feature_bc_matrix.h5 files were processed following the guidelines on the shunPykeR GitHub repository(https://github.com/kousaa/shunPykeR), an assembled pipeline of publicly available single cell analysis packages put in coherent order, that allows for data analysis in a reproducible manner and seamless usage of Python and R code. Genes that were not expressed in any cell and ribosomal and hemoglobin genes were removed from downstream analysis. Each cell was then normalized to a total library size of 10,000 reads and gene counts were log-transformed using the log(X+1) formula, in which log denotes the natural logarithm. Principal component analysis (components =20) was applied to reduce noise prior to data clustering. To select the optimal number of principal components to retain for each dataset, the knee point (eigenvalues smaller radius of curvature) was used. Leiden clustering^5^ (resolution = 0.9) was used to identify clusters within the PCA-reduced data.

Quality of the single cells (**Fig. S2A-S2B**) was computationally assessed based on total counts, number of genes, mitochondrial and ribosomal fraction per cell, with low total counts, low number of genes (≤1000) and high mitochondrial content (≥0.2) as negative indicators of cell quality. Cells characterized by more than one negative indicator were considered as “bad” quality cells. Although cells were negatively sorted prior to sequencing for the CD45 marker, a small amount of non-hematopoietic cells (expressing no Ptprc), were detected within our dataset. To remove bad quality cells and contaminants in an unbiased way, we assessed them in a cluster basis rather than individually. Leiden clusters with a “bad” quality profile and/or a high number of contaminating cells were removed. Finally, cells marked as doublets by scrublet^6^ were also filtered out. Overall, a total of 2325 cells, representing ~8.4% of all our data, was excluded from further analysis (see Figure S2 for per sample metrics). After removal of these cells, we calculated highly variable genes (HVG=3000) and re-performed PCA with unsupervised clustering analysis (components =20), followed by batch effect correction across all samples using harmony^7^ to assist annotation of cell type subsets within the dataset.

**ii) Defining Kit^lo^ and Kit^hi^ HSC subsets**

Owing to the significant drop-out effect in scRNA-seq data,^8^ it is virtually impossible to ascertain if a cell has low/mid (and sometimes high) Kit expression versus no expression. In order to methodically deal with this, we performed _data denoising and_ imputation using the MAGIC (Markov Affinity-based Graph Imputation of Cells) method,^9^ to denoise distinguish between dropouts and genuinely low gene expression values. Post-MAGIC imputation, we proceeded to set the thresholds for *Kit*, specifically, we calculated the 20^th^ and 80^th^ percentile expression value across all cells. Subsequently, cells were classified into Kit^lo^ (*Kit* expression below the 20^th^ percentile), Kit^hi^ (*Kit* expression above the 80^th^ percentile), and Kit^mid^ (*Kit* expression between the 20^th^ and 80^th^ percentile). This approach helped to enhance the quality of the data by denoising and imputing the missing values, thereby providing a more accurate representation of the gene expression landscape.

**iii) Differential expression analysis**

Differential expression analysis for comparisons of interest was performed with MAST (Model-based Analysis of Single-cell Transcriptomics) using the likelihood ratio test.^10^ In all cases, differentially expressed genes were considered statistically significant if the FDR-adjusted p-value was less than 0.05. To select maximally specific genes per cell type/subset, we ran MAST for all pairwise cluster comparisons (each versus the rest) and retained the top 10 DEGs for a given cluster (FDR<=0.05, sorted by decreasing coefficient). To generate the unique cluster signatures per se, we used the *sc.tl.score_genes*() function from *scanpy* ^11^ that calculates averaged scores based on the cluster specific genes (scores are subtracted with a randomly sampled reference gene set).

**iv) Differential abundance (DA) analysis**

DA HSC subset across age groups was identified by sampling neighborhoods of cells from a k-nearest neighbors (k-NN) graph and looking for enrichment of either age in each neighborhood as implemented in MiloR.^12^ MiloR is a graph-based statistical method to compute differential cellular abundances in neighborhoods of cells. The 15 batch-corrected latent dimensions from *scanpy* were used for MiloR (v.0.99.19) k-NN graph construction (k = 35) and neighborhood indexing (proportion = 0.1). DA testing was performed with generalized linear models, including age as covariates (neighborhoods significant if spatial corrected FDR < 0.25).

**Single- Cell ATAC sequencing**

**Preprocessing, dimensionality reduction, clustering**

Single-cell ATAC-seq from young and old mice were aligned to the mm10 genome and we processed the cellranger output file, fragments.tsv, with ArchR (v.1.0.2)^13^ was used for downstream analysis. We performed QC filtering on scATAC-seq using ArchR with the default parameters of *createArrowFiles*() (**Fig. S5**). We retained cells containing at least 1,000 and at most 100,000 fragments. Next, we filtered out the cells that did not pass QC in the corresponding scRNA-seq data, thus retaining 11493 cells common to both scRNA-seq and scATAC-seq. We then performed dimensionality reduction using iterative latent semantic indexing (LSI) on the top 25,000 variable features from the tile matrix to get a reduced dimensionality of 30 components with *addIterativeLSI*() function in ArchR. To generate visualizations, we employed the *addUMAP*() function in ArchR with the following settings: nNeighbors = 20; minDist = 0.1. For clustering, we utilized the *addClusters*() function in ArchR, specifying the parameters as follows: method = ‘Seurat’; knnAssign = 10, and maxClusters = 10. Next, we used Harmony to perform batch effect correction.

**Peak-calling and TF motif accessibility scoring**

Each individual sample was pseudobulked for peak calling with MACS2^14^ peak caller and iterative peak overlapping removal within ArchR (using default settings). We subsequently added motif annotations using *addMotifAnnotations*() with the CisBP motif database and computed chromVAR^15^ deviations for each single cell with *addDeviationsMatrix*(). To identify differentially accessible motifs within each group of interest, we applied the *rank_genes_groups*() function in *scanpy*. With the following settings: method = ‘wilcoxon’ and corr_method = ‘benjamini-hochberg’ and performed the analysis on the chromVAR zscore matrix.

**Processing Human Single-Cell CITE-seq**

We analyzed the young human bone marrow CITE-seq from Sommarin et al.^16^ using Seurat.^17^ Authors provided the peak matrix aligned to the hg38 genome. We used the sample BM_34 for which the authors provided the cell type annotations. We processed the scRNA-seq and CITE-seq data for young and old bone marrow using Seurat. The hash-tagged cells were demultiplexed using *HTODemux*() from Seurat. The cell type annotations were provided by the authors for young bone marrow samples labeled yBM1_hpc and yBM2_hpc. We predicted cell type annotations for the old bone marrow cells by label transfer using *FindTransferAnchors*() and *TransferData*() functions. We obtained the Kit^lo^ gene signature from our mouse scRNAseq data through differential gene expression analysis. To extend this signature to our human bone marrow HSPC dataset, we used Ensembl^18^ Biomart to identify human orthologues of the mouse genes. Subsequently, we scored the Kit^lo^ gene signature in human scBM HSCs using Seurat's (AddModuleScore) function.

**Data and Code Availability**

Multiome single cell RNA-and- ATAC sequencing data generated for this paper is available upon request from the corresponding author.

**Statistical Analysis**

Data were processed in GraphPad Prism 10.0 software. Statistical comparisons between 2 groups were performed with either the nonparametric unpaired Mann-Whitney U test, two-way ANOVA, ComBat batch effect correction^19^ or Wilcoxon rank sum test with continuity correction as indicated.

**Supplemental Figure Legends**

**Figure S1: (A)** Experimental schema for competitive allogeneic HCT (allo-HCT) using 2-mo (young) and 24-mo (old) 750 HSCs [Lin^−^c-Kit^+^Sca1^+^ (L^−^S^+^K^+^) CD150^+^ CD34^-^CD48^-^] from C57BL/6 mice with competitor bone marrow (BM) cells from B6.SJL-PtprcaPepcb/BoyJ mice transplanted into lethally irradiated 7-week-old (young) BALB/cJ recipients. **(B-E)** Frequency of donor-derived chimerism of all hematopoietic cells mature lineages **(B),** T-cell **(C),** Myeloid **(D),** B-cell **(E)** in the PB at the indicated time points. **(F-G)** Eight weeks after competitive HCT enumeration of absolute number of LMPP cells (LMPP/MPP4: Lineage^-^Sca-1^+^cKit^+^ Flt3^+^CD150^-^) (**F)**, CLP cells (CLP: Lineage^-^IL7Ra^+^Flt3^+^Sca^mid/lo^ Kit^lo^) **(G)**. **(H-J)** Post-HCT thymi analysis for total thymic cellularity **(H)**, enumeration of absolute number of donor-derived cells for T-cell precursors (ETP: Lineage^-^ CD4^-^ CD8^-^CD44^+^ CD25^-^Kit^+^; DN2: Lineage^-^ CD4^-^ CD8^-^CD44^+^ CD25^+^; DN3: Lineage^-^ CD4^-^ CD8^-^CD44^-^ CD25^+^) **(I)**, Mature T-cells (DP: Lineage^-^ CD4^+^ CD8^+^; SP4: Lineage^-^ CD4^+^ CD8^-^; SP8: Lineage^-^ CD4^-^ CD8^+^**) (J)**. All data are from n=9-10 mice/group. Error bars represent mean ± SEM. *P<0.05, **P<0.01, ***P<0.001, ****P<0.0001. P values calculated by nonparametric unpaired Mann-Whitney U test. Panel **A** was created using BioRender.

**Figure S2: (A)** UMAP of young HSCs after Harmony batch correction annotated by unsupervised Leiden cluster analysis (**B**), manually annotated HSC subsets (q-HSC: Quiescent HSCs; Mgk-HSCs: Platelet-biased HSCs; Mlin-HSC: Multilineage HSCs; p-HSC: Proliferative HSCs; Int-HSC: Intermediate HSCs) (**C**). (**D**) Dot plot showing marker genes for each HSC subset. Circle size and color indicate percentage and expression level, respectively. (**E**) Composition of defined HSC subtypes in Kit^hi^ and Kit^lo^ subsets in young HSCs as shown by scaled change in frequency.

(**F**) UMAP of old HSCs annotated by HSC subsets identified by *scanpy ingest*()^11^ using young HSCs, as reference dataset. **(G)** UMAP of combined young and old HSCs after Harmony batch correction annotated by sample. (**H**) Composition of defined HSC subtypes in old and young HSCs as shown by scaled change in frequency. **(I)** A neighborhood graph of the results from Milo differential abundance testing. Nodes are neighborhoods, colored by their log fold change across ages. Non-differential abundance neighborhoods (FDR 10%) are colored white, and sizes correspond to the number of cells in each neighborhood. Graph edges depict the number of cells shared between neighborhoods. The layout of nodes is determined by the position of the neighborhood index cell in the UMAP in panel **Figure 1D. (J)** Beeswarm plot of the distribution of log fold change across age in neighborhoods containing cells from different cell type clusters. Differential abundance neighborhoods at FDR 35% are colored. HSCs detected as differentially abundant annotated by HSC subsets (as defined in **Figure 1A**). **(J) (K)** Histogram for magic imputed *Kit* gene expression by age. **(L)** Representative FACS plots showing the gating strategy to define Kit^hi^ and Kit^lo^ HSC subsets in BM from young and old mice.

**Figure S3: (A-H)** Following allo-HCT (**as shown in Figure 2D**), frequency of donor-derived chimerism of hematopoietic cells **(A)**, mature lineages: Myeloid cells, B-cells, and T-cells **(B)**, in the PB at the indicated time points. **(C-D)** Enumeration of absolute number of donor-derived MP subset cells in the BM as defined in Figure S1H (**C)**, thymic non-TEC compartment (endothelial cells: CD45^-^ EpCAM^-^Ter-119 PDGFR⍺^-^ CD31^+^; fibroblasts: CD45^-^ EpCAM^-^Ter-119^-^CD31^-^ PDGFR⍺^+^) **(D)**. **(E)** Relative expression of thymic-adhesion molecules (PSGL1, CCR7, and CCR9) on donor-derived CLPs (Statistical analysis was performed using ComBat^19^ to perform batch-effect correction for the mean MFI across two experiments), **(F)** Absolute amount of intrathymic thymopoietic ligands, IL-22 (left), RANKL (right) was measured by ELISA after allo-HCT. **(G-H)** Frequency of donor-derived **(G)** CD4+ and **(H)** CD8+ T-cell (naïve: CD62L+CD44- ; Central Memory: CD62L+CD44+; Effector Memory: CD62L-CD44+) subsets in the spleen. All data are from n=10-11 mice/group. **(I-O)** Following allo-HCT (**as shown in Figure 2P**)**,** frequency of donor-derived chimerism of mature lineages, myeloid (**I**) and B-cells (**J**) in the PB at the indicated timepoints. **(K-O)** Enumeration of absolute number of donor-derived LSK **(K) and** MP subset cells (**M**) in the BM as defined in Figures 2F and S1H, respectively**,** thymic TEC **(N) and** non-TEC **(O)** compartments as defined in Figures 2K and S3D**.** Analyses were performed at 16 weeks following competitive HCT. All data are from n=9-10 mice/group.

**Figure S4:** Competitive allo-HCT using Kit subsets from young mice, as described in Figure 2D. Twenty weeks after competitive HCT, (**A**) frequency of donor-derived chimerism of mature lineages in the peripheral blood as described in Figure 2E.Enumeration of absolute number of donor-derived cells in the BM for LSK cell subsets as defined in Figure 2F (**B**), MP cell subsets as defined in Figure S1H (**C**), CLP cells as defined in Figure 2G (**D**), total thymic cellularity (**E**), donor-derived mature T-cells (**F**) and T-cell precursor thymocyte subsets (**G**) as defined in Figures 3I-3J, thymic stromal compartment (CD45- cells) as defined in Figures 2K and S3D (**H**). All data are from n=8 mice/group. Error bars represent mean ± SEM. *P<0.05, **P<0.01, ***P<0.001. P values calculated by nonparametric unpaired Mann-Whitney U test.

**Figure S5: (A-E)** Single-cell ATAC seq pre-processing and QC analysis. **(A)** Comparison of TSS enrichment score and number of fragments. The dashed lines depict the cutoffs used. **(B)** Ridge plots showing distribution of TSS enrichment scores per sample (**top**) and distribution of unique nuclear fragments per sample (**bottom**). **(C)** Plots showing fragment size distribution (**top**) and TSS enrichment profile (**bottom**) across samples. (**D**) scATAC-seq UMAP without batch correction colored by sample. **(E)** Volcano plots of differentially accessible (DA) peaks (fold change> 2, FDR <0.01) for HSC subset Cluster 3 vs. Cluster 6 (Wilcoxon rank-sum test).

**Figure S6:** Zbtb1 expression assessed by flow cytometric analysis following knock out (KO) in Rosa26Cas9 KI Kit^lo^ HSCs **(A)** and OE of Zbtb1 cDNA in Kit^hi^ HSCs **(B)**.

**Figure S7**: (**A-F**) Following allo-HCT (**as shown in Figure 3C**)**,** frequency of donor-derived chimerism of mature lineages in the PB **(A)**. **(B-F)** Enumeration of absolute number of donor-derived MP subset cells in the BM as defined in Figure S1H **(B),** donor-derived mature T-cells as defined above **(C)**, thymic non-TEC compartment as defined in Figure S3D **(D),** and donor-derived **(E)** CD4+ T-cell and **(F)** CD8+ T-cell subsets in the spleen as defined above. All data are from n=9-10 mice/group. All analyses were performed at 8 weeks following competitive HCT. **(G)** Following allo-HCT (**as shown in Figure 3L**), frequency of donor-derived chimerism of mature lineages in the peripheral blood. Analyses were performed at 8 weeks following allo-HCT. All data are from n=5-6 mice/group. **(H-O)** Competitive allo-HCT using Kit subsets from old mice, as described in Figure 3C. Twenty weeks after competitive HCT, (**H**) frequency of donor-derived chimerism of mature lineages in the peripheral blood as described in Figure 2E.Enumeration of absolute number of donor-derived cells in the BM for LSK cell subsets as defined in Figure 2F (**I**), MP cell subsets as defined in Figure S1H (**J**), CLP cells as defined in Figure 2G (**K**), total thymic cellularity (**L**), donor-derived mature T-cells (**M**) and T-cell precursor thymocyte subsets (**N**) as defined in Figures 3I-3J, thymic stromal compartment (CD45- cells) as defined in Figures 2K and S3D (**O**). All data are from n=6-7 mice/group. Error bars represent mean ± SEM. Error bars represent mean ± SEM. *P<0.05, **P<0.01, ***P<0.001, ****P<0.0001. P values calculated by nonparametric unpaired Mann-Whitney U test.

**Figure S8:** Equal Competition allo-HCT using Kit^lo^ HSCs from young and old mice. (**A**) Experimental schema for equal competition allogeneic HCT (allo-HCT) using Kit^lo^ HSCs from 2-mo young (blue) and 22-24-mo old (light-blue) C57BL/6 mice combined with competitor bone marrow (BM) cells from B6.SJL-PtprcaPepcb/BoyJ mice transplanted into lethally irradiated 7-week-old (young) BALB/cJ recipients. Twenty weeks after competitive HCT, (**B**) frequency of donor-derived chimerism of mature lineages in the peripheral blood as described in Figure 3B. Enumeration of absolute number of donor-derived cells in the BM for LSK cell subsets as defined in Figure 2F (**C**), MP cell subsets as defined in Figure S1H (**D**), CLP cells as defined in Figure 2G (**E**), donor-derived T-cell precursor thymocyte subsets as defined in Figure 3I (**F**) and mature T-cells as defined in Figure 3J (**G**). All data are from n=9 mice. Error bars represent mean ± SEM. (**H**) Pathway enrichment analysis performed on differentially expressed genes comparing old versus young Kit^lo^ HSCs by *gseapy*()^20^ using Hallmark_2020 and GO_Biological_Process_2023 libraries. Bubbleplot showing representative pathways enriched in young Kit^lo^ HSCs. (**I-K**) Matrix plots showing motif enrichment identified by ChromVAR (I), gene accessibility score (**J**), and gene expression (**K**) for Kit^lo^ subset in young and old HSCs. *P<0.05, **P<0.01, ***P<0.001. P values calculated by nonparametric unpaired Mann-Whitney U test.

**Figure S9: (A**) UMAP of CD34+ young and old human BM CITE-seq^16^ annotated with 15 clusters. **(B)** Violin plot for CD117 ADT-expression by age. Statistical analysis performed using Wilcoxon test. **(C)** Representative FACS plots of human BM samples showing gating strategy to define HSPC subsets. **(D)** Representative FACS plots of human BM samples showing gating strategy to define KIT^hi^ and KIT^lo^ HSCs in human BM.

**Supplemental Data Table**

**Data Table 1:** Top marker genes for each HSC subset.

**Data Table 2:** Dataset of differentially expressed genes (DEGs) comparing young and old HSCs at steady state.

**Data Table 3:** Zbtb1-targets enriched in Kit^lo^ HSCs

**Data Table 4:** Pathway enrichment analysis of Non-Notch1 Zbtb1 targets enriched in Kit^lo^ HSCs

**Data Table 5:** Pathway enrichment analysis of differentially expressed genes comparing young and old Kit^lo^ HSCs

**Data Table 6:** Human Bone Marrow Donor Information
